# Interchromosomal Linkage Disequilibrium Analysis Reveals Strong Indications of Sign Epistasis in Wheat Breeding Families

**DOI:** 10.1101/2025.05.16.654524

**Authors:** Augusto Tessele, Geoffrey Morris, Eduard Akhunov, Blaine Johnson, Marshall Clinesmith, Addison Carroll Hope, Allan Fritz

## Abstract

Additive gene action is assumed to underly quantitative traits, but the eventual poor performance of elite wheat lines as parents suggests that epistasis could be the underlying genetic architecture. Sign epistasis is characterized by alleles having either a beneficial or detrimental effect depending on the genetic background, which can result in elite lines that fail as parents in certain parental combinations. Hence, the objective of this study were to test the existence of sign epistasis and examine its consequences to wheat breeding. The presence of sign epistasis is expected to distort the allele frequency distribution between two interacting genes compared to neutral sites, creating strong linkage disequilibrium (LD). To test this hypothesis, analysis of interchromosomal LD in breeding families was performed and detected 19 regions in strong disequilibrium, whose allele frequency distribution matched the sign epistasis prediction and falsified the competing hypothesis of additive selection. To validate these candidate interactions while avoiding the biases of a circular analysis and the confounding effects of genetic drift, two independent sets of populations were analyzed. Genetic drift was attributed to creating the sign epistasis patterns observed in eleven interactions, but there was not sufficient evidence to reject the sign epistasis hypothesis in eight interactions. Sign epistasis may explain the poor performance of elite lines as parents, as crossing lines with complementary allelic combination re-establishes epistatic variance in the offspring. Reduction in the effective population size in certain crosses may also occur when unfavorable sign epistatic combinations are deleterious. The potential existence of di-genic and higher order epistatic interactions in elite germplasm can tremendously impact breeding strategies as managing epistasis becomes imperative for success.

## Introduction

The infinitesimal model (Fisher, 1918) proposes that additive effects play a primary role in genetic inheritance, while relegating dominance and epistasis to lesser importance (Turelli, 2017). However, the eventual occurrence of elite lines performing poorly as parents suggests that epistasis might be the underlying genetic architecture. Sign epistasis, a significant form of gene action in evolutionary genetics (Wright, 1932), is characterized by alleles exhibiting either advantageous or detrimental effects depending on the allelic state of the interacting gene (Weinreich et al., 2005; Poelwijk et al., 2007). Under the influence of sign epistasis, genotypes can adapt to environmental pressures in various combinations of alleles, often leading to similar adaptation (Wright, 1988). In breeding, it implies the existence of elite genotypes with different frameworks of favorable allelic combinations. Hence, crossing contrasting genotypes causes the favorable allelic combinations to segregate in the offspring and results in mean performance lower than the midparent value. This clash of epistatic interactions between parental lines may justify the eventual poor performance of elite lines as parents. For instance, the widely grown hard red winter wheat cultivars *Jagger* and *2137*, both released in 1994, had comparable agronomic performance, reaching 34.8% and 23.1% of the acres in Kansas in 2000, respectively, yet *Jagger* appears in the pedigree of 21 subsequent releases, whereas *2137* contributed to only one (USDA ARS n.d.).

Establishing the importance and extent of epistasis in wheat breeding by modeling epistatic variance has tremendous challenges because of the lack of orthogonality with additive effects (Tessele et al., 2024; Raffo et al., 2022; Vitezica et al., 2017). However, the existence of sign epistasis interactions under selection may leave detectable signatures in the genome. The influence of sign epistasis in each locus closely resembles the effects of balancing selection, which maintains genetic diversity at sites under selection and can augment diversity at linked loci (Lewontin and Hubby, 1966; Hudson and Kaplan, 1988; Charlesworth et al., 1997; Takahata and Satta, 1998). The polymorphism flanking the loci under balancing selection can span over large or short regions, depending on local recombination frequency and strength of LD (Charlesworth et al., 1997; Wiuf et al. 2004; Charlesworth, 2006), the age of the advantageous polymorphism and the selection intensity (Kreitman and Rienzo, 2004; Tian et al., 2002). Besides affecting each locus independently, sign epistasis also impacts the relationship between interacting genes because selection acts on allelic combinations rather than on individual alleles. In a two-way interaction, the preferential selection of favorable allelic combinations distorts the allele frequency distribution of interacting genes compared to two neutral sites, creating strong disequilibrium that can be regarded as a selection signature.

Although selection for sign epistasis may leave signatures in the genome, the mere existence of LD does not imply sign epistasis. For instance, short-range linkage disequilibrium (SRLD) primarily arises due to random genetic drift or common ancestry of chromosomal blocks (Koch et al., 2013). SRLD are perpetuated under low recombination, making it virtually impossible to disentangle from an eventual sign epistasis signature. Long-range linkage disequilibrium, however, suggests that a countervailing force to recombination maintains strong LD between distant sites, which is expected to be even more prominent in interchromosomal sites because the independent assortment of chromosomes recombines physically unlinked loci in every meiotic event. If sign epistasis is pervasive in wheat, breeding programs may perpetuate interchromosomal LD by continuously selecting favorable interactions. Detecting true selection signatures of epistasis (Lewontin et al., 1960) is challenging as other confounding effects can also create long-range LD, as population admixture (Maccaferri et al., 2005; Chao et al., 2010), genetic drift (Koch et al., 2013), artificial selection (Flint-Garcia et al., 2005; Joukhadar et al., 2019), or structural variation in chromosomes (Zhao et al., 2022). Disentangling signatures of epistasis from genetic artifacts may be possible through examining the allele frequency of sites in strong LD, as additive selection is expected to create a different allele frequency distribution than epistasis. Population structure can be minimized by analyzing breeding families independently. Analyzing interactions independently from the base population used for discovery may reduce the bias steaming from genetic drift and prevent statistical inference on a circular analysis (Kriegeskorte et al., 2009).

The main hypothesis is that sign epistasis exists and has major implications to wheat breeding. The objectives were to: (i) validate the existence of loci in abnormally strong interchromosomal linkage disequilibrium; (ii) test if selection for sign epistasis creates the observed strong interchromosomal LD; (iii) verify the existence of and disentangle confounding genetic artifacts that could likewise create strong interchromosomal LD.

## Material and Methods

### Plant Material and Genomic Data

The dataset consisted of genomic information on experimental lines from the Kansas State Hard Red Winter Wheat breeding program for the years 2011 through 2021, which were categorized based on the last stage of the breeding pipeline reached by each experimental line. The four stages in the breeding pipeline were early, preliminary, advanced, and elite yield trials, named IPSR, PYN, AYN and EYN, respectively. All experimental lines were genotyped in the IPSR stage using genotype-by-sequencing (GBS) (Poland et al. 2012). As part of the genotyping pipeline of the K-State wheat breeding program, markers with a MAF <0.01, more than 20% missing values and more than 20% of heterozygotes were filtered, as described in Jarquin et al (2017). The R package *‘*rrBLUP *‘*(Endelman, 2011) was used for marker imputation, where missing markers were imputed with the mean value among all lines for that marker and resulted in a total of 55,148 SNPs.

### Validating the existence of strong interchromosomal linkage disequilibrium

This exploratory stages sought to validate the existence of interchromosomal linkage disequilibrium in breeding germplasm. To test this hypothesis, experimental lines from the EYN stage of the breeding program from 2010 to 2021 were considered. A linkage disequilibrium analysis (r^2^) focusing only on interchromosomal interactions was conducted and identified pairwise markers in strong linkage disequilibrium. Validating the existence of strong interchromosomal LD in wheat enabled further investigation into the hypothesis of selection signatures of sign epistasis (Figure 1).

**Figure 1.**
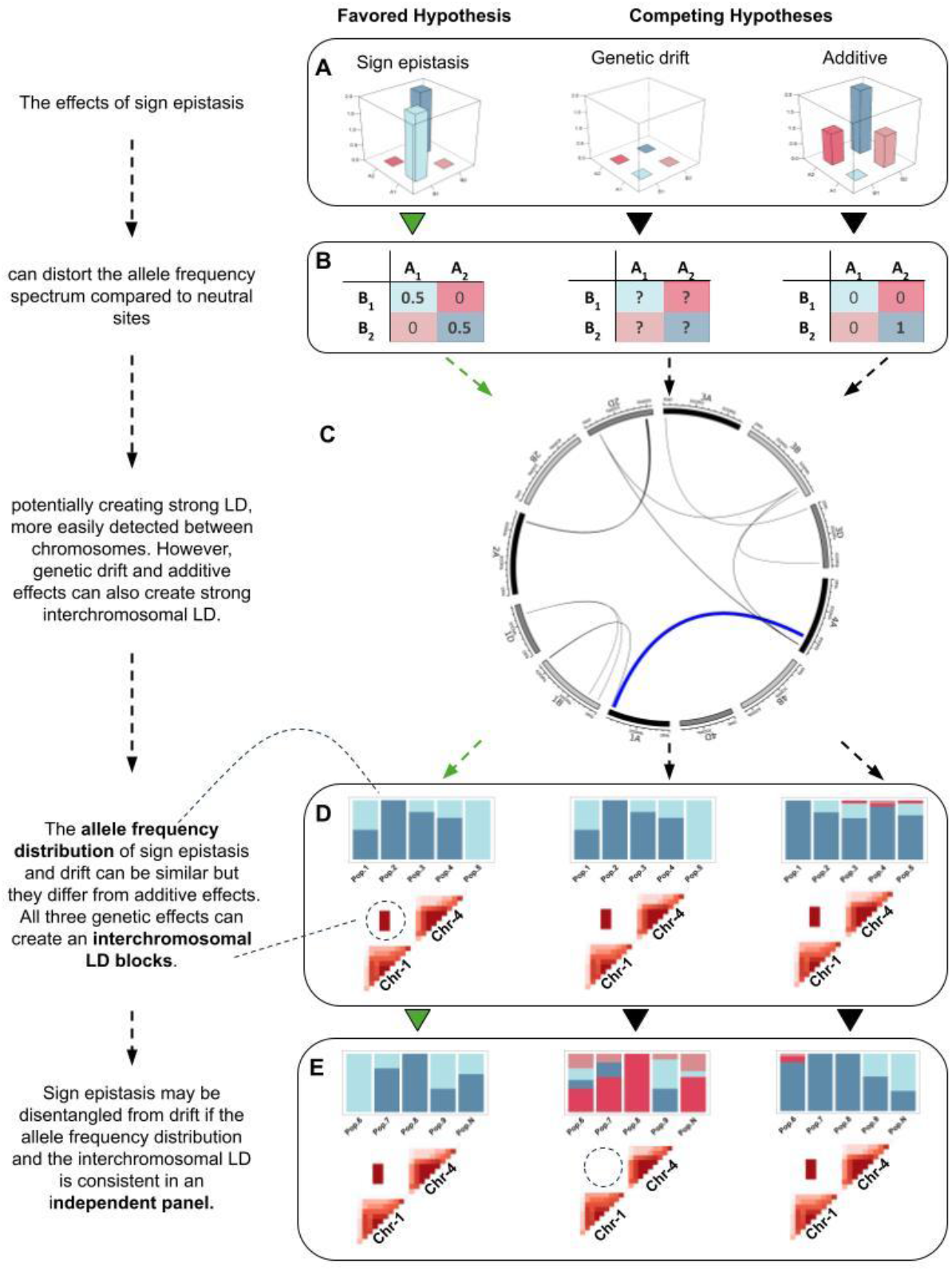
Selection of sign epistasis interactions can create interchromosomal LD. **A** depicts the levels of adaptation of two genes (A and B) under the effects of sign epistasis, drift and additive. **B** shows the expected allele frequency distribution under an ideal scenario for each genetic effect. **C** illustrates strong linkage disequilibrium between pairwise interchromosomal sites, where inner links connect sites in strong LD. **D** represents the allele frequency distribution in five populations for two genes in strong LD and the respective heatmap of intra- and inter-chromosomal LD for each genetic effect. **E** represents the allele frequency and LD values in an independent panel from D, which highlights the consistency of sign epistasis contrasting to the randomness of genetic drift.

### Minimizing the influence of population structure and genetic drift

Historical data from 25 breeding families was used for the discovery of selection signatures indicative of sign epistasis (Figure 1C). These were large families tested in the early yield trial stage of the breeding pipeline, with sizes ranging from 20 to 45 genotypes. As a data quality measure, a principal component analysis was conducted for each of the families to identify and exclude any outlier genotypes detected in the two-dimensional plot defined by the first two principal components to mitigate the potential introduction of population structure resulting from genotyping or seed processing errors.

An interchromosomal linkage disequilibrium (r^2^) analysis was conducted only on markers with MAF>0.3 to reduce the total number of interactions to screen for and to increase the likelihood of detecting interactions with allele frequency characteristic of sign epistasis (Figure 1A and B). To circumvent the effect of population structure, the interchromosomal LD analysis was conducted in each family individually and all interactions presenting r^2^ > 0.9 were retained. Although this approach controls population structure, it was highly influenced by genetic drift within families. To address the genetic drift effect, the combined data of the 25 families was used to run another interchromosomal LD analysis, and all interactions presenting r^2^>0.8 were selected. Interactions selected in both analyses and observed in a minimum of two families were deemed worthy of further investigation. Supplemental Table 2 demonstrates this theoretical framework for this approach.

### The Balancing Selection Nature of Sign Epistasis and Interchromosomal Linkage Blocks

A sign epistatic interaction under selection, besides the characteristic intrachromosomal haplotype block, may also create an “interchromosomal block” (Figure 1D). To investigate this hypothesis, 40 markers flanking each candidate sign epistatic marker were used to run a linkage disequilibrium analysis, and the resulting intra and interchromosomal LD values were used to construct heatmaps of 2D LD plots.

### Allelic Frequencies to Differentiate Additive and Sign Epistasis

Selection for sign epistasis is expected to change allele frequencies differently than additive selection (Figure 1B). To empirically test this hypothesis, the allele frequencies of each candidate interaction was computed, and the patterns were analyzed (Figure 1D). Since these families were genotyped in the F_5:6_ generation using GBS, heterozygous loci were still observed.

### Independent Family and Breeding Program Validation

To minimize the effects of genetic drift and to avoid a circular analysis, 15 distinct validation families were selected to reanalyze the heatmaps of 2D LD plots and the distribution of allele frequencies in order to validate candidate interactions (Figure 1E). Although independent from the discovery set, the total number of families analyzed was relatively small and prone to the effects of genetic drift. To diminish the confounding effects of drift in small populations, data from a time span of 10 years in the wheat breeding program was also utilized to further validate the consistency of candidate interactions. The breeding program dataset was divided sequentially from early to elite yield trials and allele frequencies were calculated for each stage. In addition, the combined data of the breeding program was used to falsify candidate interactions if the opposite combination (relative to the most common allelic combination) had less than 10% of the total allelic combinations.

### Simulation to Estimate Theoretical Distribution of Interchromosomal LD Values

To evaluate interchromosomal linkage disequilibrium (LD) attributable solely to genetic drift, we simulated 100 breeding crosses using random lines from the Kansas State University wheat breeding program as parents. Both single- and three-way crosses were generated, and the resulting populations (15 individuals each) were selfed to the fifth generation. Interchromosomal LD was assessed within each family individually and across all families combined. The same thresholds used to detect candidate interactions in the empirical discovery families were also applied to the simulated families and the results were compared with those from empirical discovery families. Simulations were performed using the R package AlphaRSim (Faux et al., 2016).

### Software and Packages

The linkage disequilibrium analysis was performed using the software PLINK (Purcell et al., 2007). The R software packages *“*factoextra” (Kassambara et al., 2017), *“*circlize” (Gu and Gu, 2022), and *“*gaston” were utilized to perform the principal component analysis, to plot the SNPs presenting strong interchromosomal LD, and to plot the heatmap of 2D LD plots, respectively. The package “ggplot2” was used to plot allele frequency counts (Wickham, 2016). In addition, built-in functions in R were used during data management and to calculate allele frequencies.

## Results

### Simulated versus empirical interchromosomal linkage disequilibrium

The simulated families, generated under the sole effects of random genetic drift, displayed numerous pairwise interactions exhibiting some degree of interchromosomal linkage disequilibrium (LD) in at least two families, but only a few with R^2^ values approaching 0.9 (Figure 2). In contrast, the empirical discovery families exhibited a much larger number of pairwise interactions exceeding these thresholds (Figure 2).

**Figure 2.**
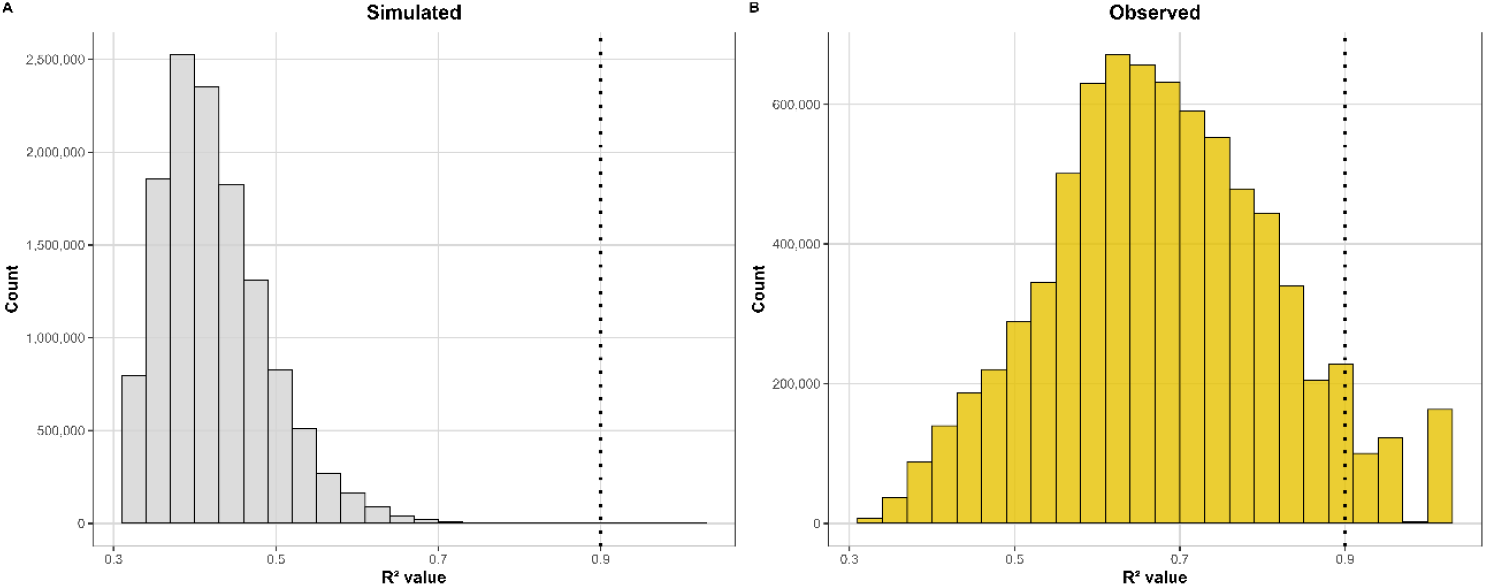
Simulated and observed distribution of interchromosomal R^2^ values found in at least two families. (A) Simulated distribution of interchromosomal LD R^2^ values under the sole influence of genetic drift. (B) Observed distribution of interchromosomal LD R^2^ values in empirical discovery panel. The vertical dashed line marks the first threshold used to select pairwise interactions for further analysis.

When marker data from all populations were combined, the simulated families showed very few interactions with R^2^ values near or above 0.8 (Figure 3), whereas the discovery families displayed a far greater number of pairwise interactions surpassing this threshold. Importantly, none of the simulated interactions met the criteria used to designate candidate interactions— namely, an average R^2^ > 0.9 in at least two families individually and R^2^ > 0.8 when analyzed across all populations combined.

**Figure 3.**
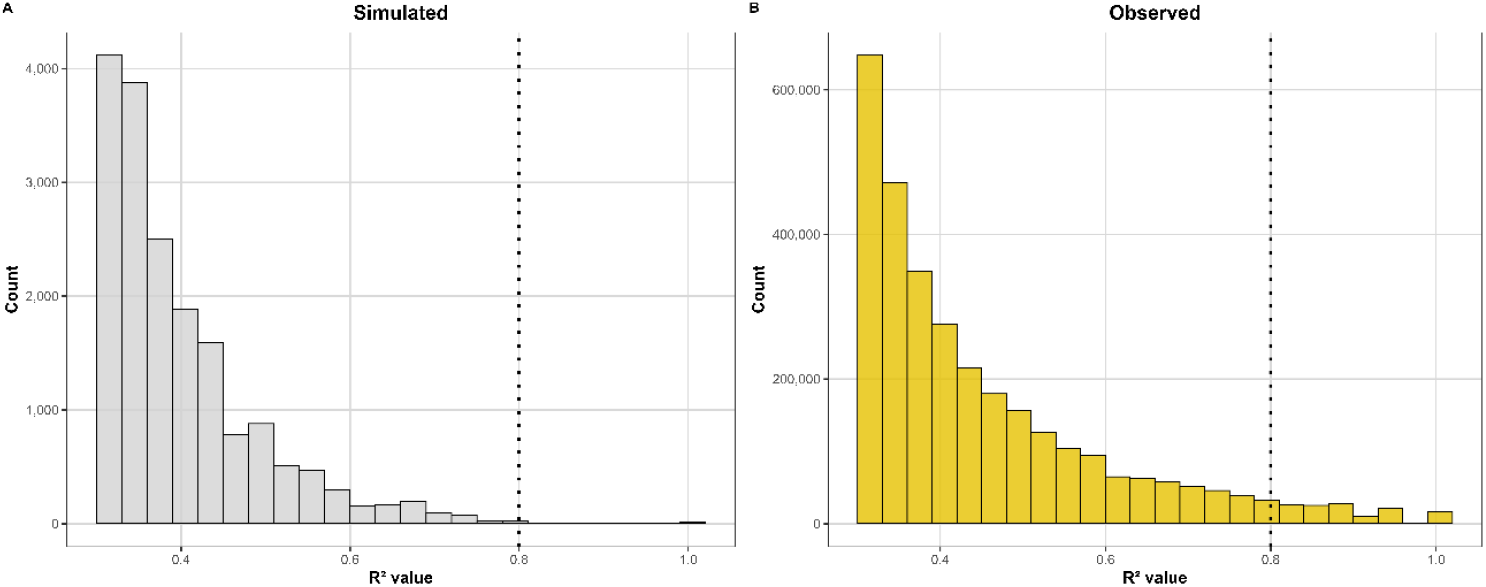
Simulated and observed distribution of interchromosomal R^2^ values when analyzing populations collectively. (A) Simulated distribution of interchromosomal LD R^2^ values under the sole influence of genetic drift. (B) Observed distribution of interchromosomal LD R^2^ values in empirical discovery panel. The vertical dashed line marks the second threshold used to select pairwise interactions for further examination.

### Detection of loci in strong interchromosomal linkage disequilibrium

The goal of the discovery stage was to pinpoint a manageable number of interactions in strong LD while minimizing the likelihood of identifying artifacts of population structure, genetic drift, or additive selection. In total, 269 pairwise SNPs, which represent 19 interacting genomic regions, passed the set thresholds, and were considered candidate selection signatures (Figure 4). Wheat subgenome B had the most candidate interactions with 15, followed by A and D with 13 and 10, respectively. The chromosomes with the most candidate interactions were 1B and 6B, with four interactions each. In total, 14 out of the 19 interactions were intergenomic, and most involved subgenomes B and D. Subgenome A and B had candidate intragenomic interactions, which were absent in subgenome D. In addition, a candidate of a three-way selection signature was also observed in chromosomes 2D, 3B and 4A (Figure 4). A total of seven candidate interactions had opposed allelic combinations fixed in different families, and 12 were fixed for one allelic combination while the opposed allelic combination was observed in segregating families (Supplemental Table 1).

**Figure 4.**
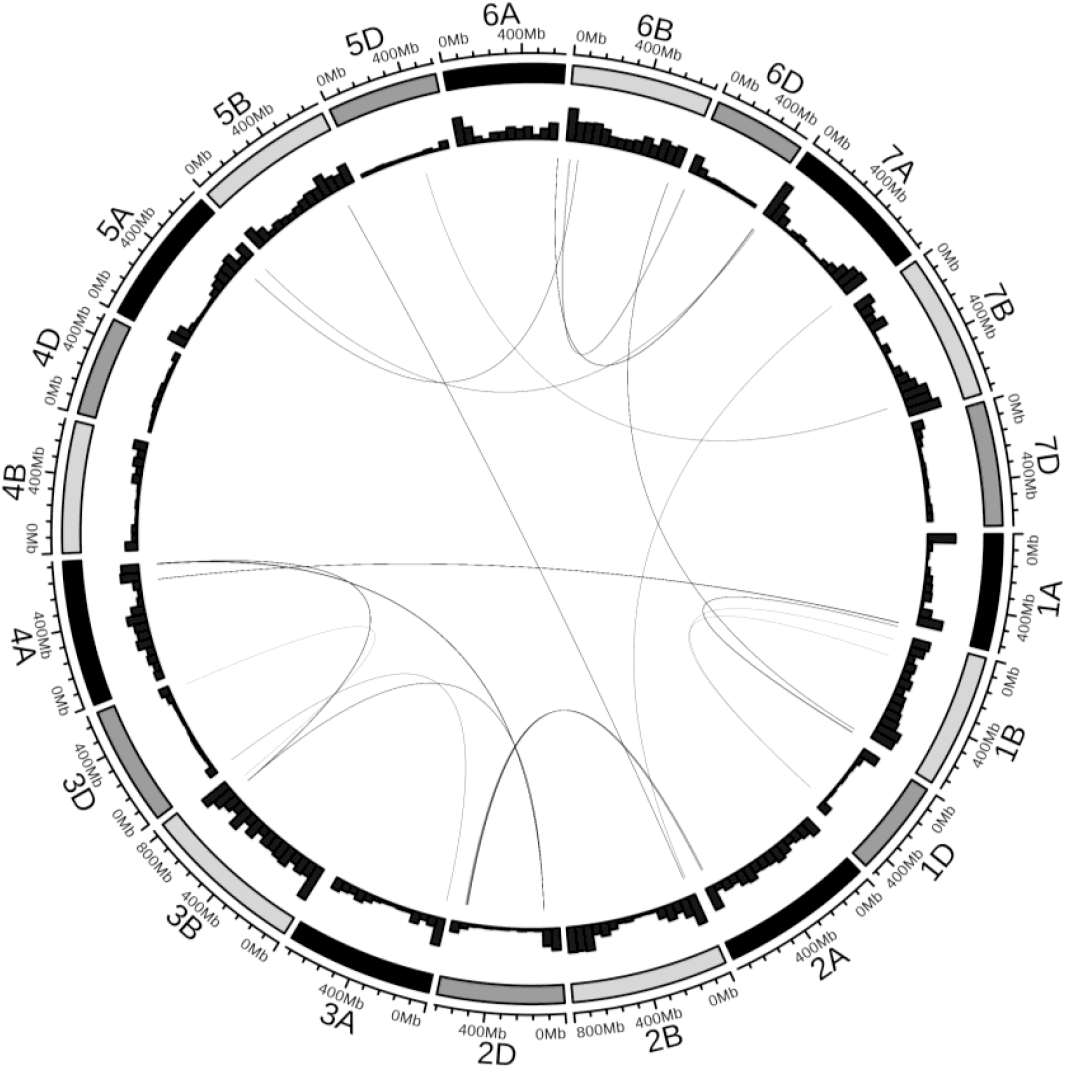
Candidate selection signatures of sign epistasis. The inner links represent regions (SNPs) in strong interchromosomal LD following the discovery approach previously described. The outer histogram plots represent the marker density in each chromosome. The outermost bar represents the 7 chromosomes and 3 subgenomes of wheat.

### Putative balancing selection associated with sign epistasis

The analysis of linkage drag was performed for each of the 19 candidate selection signatures of sign epistasis. The results for the candidate interaction involving chromosomes 1A and 4A reveal the presence of “interchromosomal linkage blocks” associated with the candidate selection signature (Figure 5B). These blocks are primarily attributed to a sizable linkage block located on chromosome 1A and a smaller block situated on chromosome 4A. The control interaction, however, displayed some level of intrachromosomal LD on chromosome 2B but no “interchromosomal LD block” was observed (Figure 5B).

**Figure 5.**
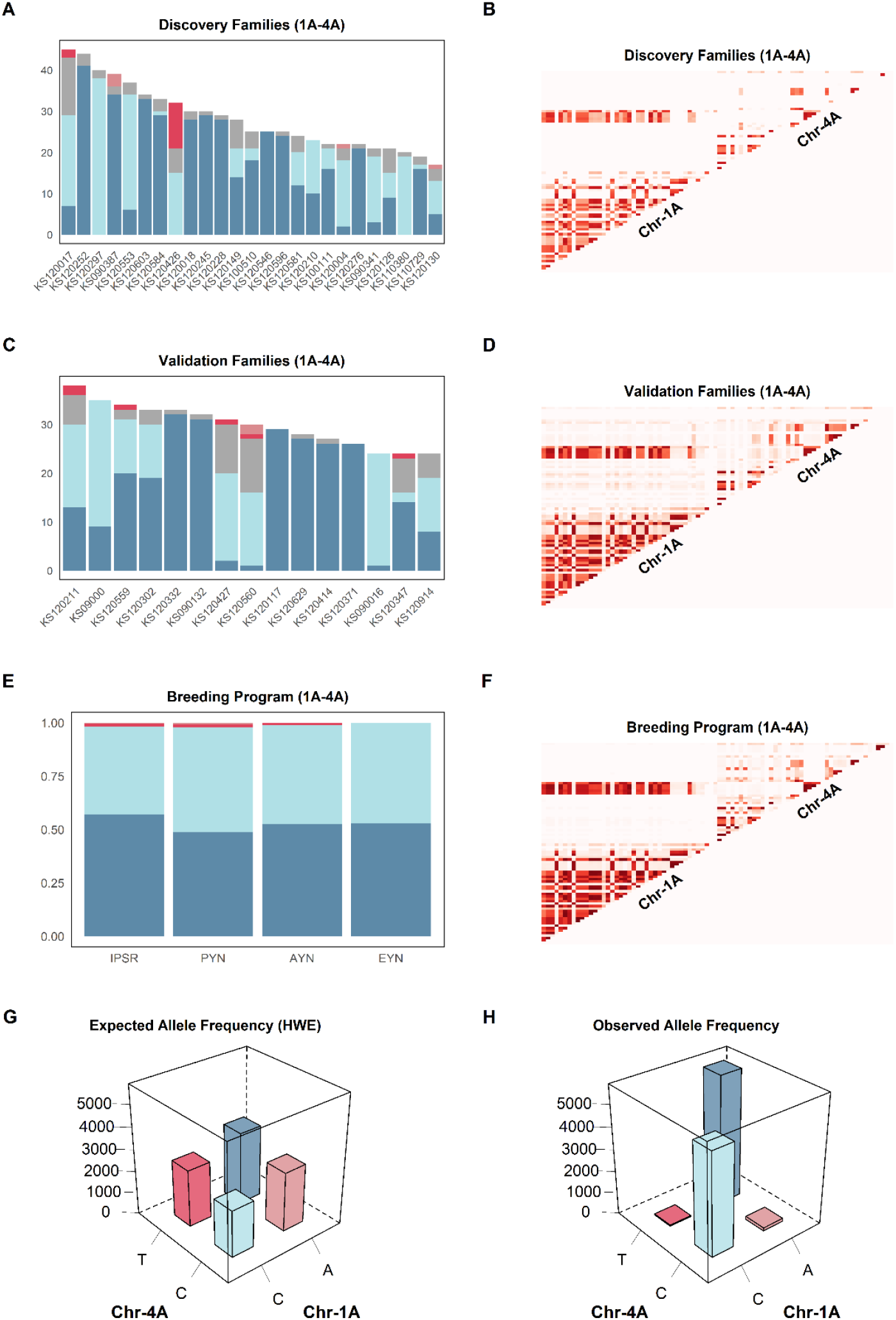
Testing independent breeding panels for sign epistasis. Allele frequency distribution (left column) and heatmaps of 2D LD plots (right column) of the candidate epistatic interaction involving segments of chromosomes 1A and 4A. **A, C** and **E** display the allele frequency distribution from the discovery and validation panels, and from the last four stages of the breeding pipeline, respectively. **B, D** and **F** shows the 2D LD plot from discovery and validation panels, and from the last four stages of the breeding pipeline, respectively. **G** displays the expected allele frequency distribution in the entire breeding program under Hardy-Weinberg Equilibrium and **H** shows the observed allele frequency distribution.

The remaining interactions presented *“*interchromosomal linkage blocks” of variable sizes, as the number of flanking SNPs in strong LD with the candidate interaction was variable (Supplemental Figures 1-18). For example, candidate interactions 5D-7B, 2D-3B, 1BS-1D and 1BL-1D were in intrachromosomal LD with a few SNPs, resulting in very small *“*interchromosomal LD blocks”.

### Allele frequencies disentangle epistasis from additive selection

The allele frequencies of candidate interaction 1A-4A in the discovery stage reveals many families nearly fixed for the A/T and C/C allelic combinations (Figure 5A). Families segregating for these two alleles exhibited a high frequency of opposed allelic combinations, suggesting that selection is actively favoring both the A/T and C/C allelic combinations. Off-combinations consistently presented very low frequency in most families. The absence of a predominant allelic combination in high frequency among segregating populations weakens the competing hypothesis of additive selection.

The candidate interactions 2B-5B, 3B-3D, and 6B-6Da had a similar distribution of opposing allelic combinations as interaction 1A-4A (Supplemental Figures 5:7-A). Candidate interactions 2D-4A, 2A-2D, 2D-3B, 3B-4A and 5A-7A had a common pattern where one of the contrasting combinations predominated in most families, followed by the opposed combination as the second most frequent, and followed by heterozygotes (Supplemental Figures 1:4-A, 14-A). However, for interactions 1B-6B, 5D-7B, 3A-3D, 1A-1B, 5A-6B, 6A-7A, 1BL-1D, 1BS-1D, 2B-7A and 6BL-6D, the most common allelic combination was found at a very high frequency in most families, and the opposing combination was observed in only a few families (Supplemental Figures 8:13-A, 15:18-A). In all cases, the off-combinations were observed at very low frequencies or absent in many families.

The allele frequencies of the candidate three-way epistatic interaction indicates the predominance of the C/C/C alleles, which was fixed in many families (Supplemental Figure 19-A). In segregating families, the opposing allelic combination (A/T/T) was found in high frequency, suggesting that both combinations could be under positive selection. Off-combinations were less common than heterozygotes in most families.

### Validation Families Support Sign Epistasis Hypothesis

The patterns of LD in the validation families remained consistent with the discovery families for interaction 1A-4A, where a long linkage block on chromosome 1A and a small block on 4A created the characteristic *“*interchromosomal linkage block” (Figure 5-D). The frequency of the allelic combinations in the validation families presented a similar pattern as the discovery families (Figure 5-C). Six populations are fixed for the A/T combination, and the remaining 11 are segregating with both A/T and C/C as the most frequent combinations. The frequency of off-combinations was also very low.

Interaction 2B-5B (Figure 5-C) had a similar pattern of allelic combination as interaction 1A-4A, where the validation families had an allele frequency pattern similar to the discovery families. The interchromosomal linkage analysis for interactions 1B-6B, and 5D-7B did not present the characteristic *“*interchromosomal LD block” (Supplemental Figure 8-D and 9-D) because most families were fixed for the major allelic combination (Supplemental Figure 8-C and 9-D).

On the allele frequency analysis, the candidate interactions 5A-7A, 1BL-1D, 1BS-1D, 2B-7A and 6B-6Db (Supplemental Figures 14:18-C) had at least one family with an off-combination as the most frequent allelic combination. This suggests that these loci are not under selection, and the sign epistasis hypothesis was rejected in these candidate interactions. For interaction 6B-6D*a*, family KS120427 had an off-combination as the most frequent allelic combination. However, the abnormal high frequency of heterozygotes, which could have a biological meaning or simply be the consequence of genotyping errors, prevents drawing any definitive conclusions about the interaction (Supplemental Figure 7-C). Similarly, for interaction 3B-3D, the family KS090132 was fixed for an off-combination but the remaining families presented allelic frequencies matching the sign epistasis hypothesis (Supplemental Figure 6-C). It is probable that the parental lines of population KS090132 were fixed for the off-combination, which prevents observing the favorable candidate combinations. Thus, this interaction was kept for further analysis.

**Figure 6.**
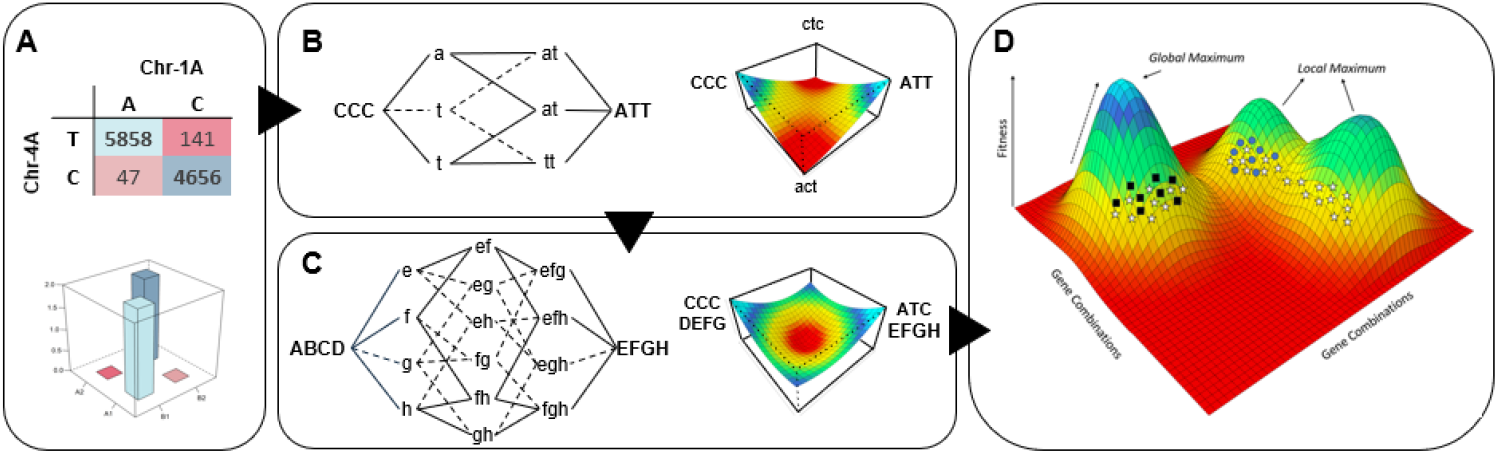
Rugged fitness landscapes are formed from sign epistasis interactions. **A** The allele frequency of interaction 1A-4A across the breeding program indicates a di-genic sign epistasis interaction. **B** The potential three-gene sign epistasis interaction involving chromosomes 2D, 3B and 4A creates a valley of adaptation between the two optimum allelic combinations. **C** Combining a hypothetical four- and the potential three-gene sign epistasis interaction creates a fitness landscape with regions of high, intermediate, and low adaptation. Factual representation of the fitness landscape is difficult because of the elevated complexity of the genetic system. **D** Rugged fitness landscape formed by several sign epistasis interactions with variable levels of gene interactions. This is an extreme simplification of a highly complex, multi-dimensional genetic system. Adaptations from Wight (1932).

For the remaining interactions, a consistent pattern emerged where one allelic combination predominated in the majority of families, either fixed or at a very high frequency, while the opposing allelic combination was observed in a few other families at varying frequencies (Supplemental Figure 1:4-C, 10:13-C). The candidate three-way interaction was segregating in few populations, with the allelic combination C/C/C as the most frequent, followed by the opposed allelic combination (A/T/T) and a small occurrence of off-combinations (Supplemental Figure 19-C). The remaining families were fixed for the C/C/C allelic combination.

### Breeding Program Validation

To further test if the 14 non-falsified interactions are not the result of strong genetic drift effects in small population sizes, data from 10 years of the breeding program was used to calculate the frequency of allelic combinations. The results for interaction 1A-4A (Figure 5-E) reveal that off-combinations were present at low frequencies in the early stages of yield trials (IPSR). The frequency of off-combination declined with successive generations of selection, ultimately leading to complete elimination of off-combinations in the EYN.

The same pattern was observed for interactions 6B-6Da, 5A-6B, and 6A-7A (Supplemental Fig 7-E, 12-E, and 13-E), which exhibited no or very few off-combinations at the EYN stage. However, the latter two candidate interactions had the opposite combination with less than 10% of the total allelic combinations, suggesting that the detection of these interactions may have arisen due to genetic drift. Consequently, the sign epistasis hypothesis for the 5A-6B and 6A-7A interactions was rejected. Similar allele frequencies were observed for interactions 3A-3D and 1A-1B, leading to the rejection of these combinations as well (Supplemental Figure 10-E and 11-E). Interactions 1B-6B and 5D-7B presented one off-combination with similar frequency than the favored, opposed allelic combination and were also discarded (Supplemental Figure 8-E and 9-E).

The six remaining interactions (2A-2D, 2D-4A, 2D-3B, 3B-4A, 2B-5B, and 3B-3D) displayed a high frequency of both opposing allelic combinations but showed minor variations in off-combinations across the breeding stages (Supplemental Figures 1:6-E) and were not falsified due to allele frequency. However, interactions 2D-3B, 3B-4A, 2B-5B, 3B-3D, and 6B-6Da had at least one of the two interacting regions involving a single marker, where a single marker alignment error in the GBS pipeline could result in the observed LD and allele frequencies. For such reason, these interactions were discarded. There was not enough evidence to discard interactions 2D-4A and 2A-2D.

Interactions 3B-3D, 2B-5B, and 2A-2D had an average of 94% of genotypes with the two favorable contrasting allelic combinations, at an average ratio of 4:1 between the most common allelic combination and its opposed combination. The interactions 2D-3B, 2D-4A, and 3B-4A comprised the candidate three-way sign epistasis interaction and had the two favorable interactions in very high frequencies across the four breeding stages (Figure 19-E). The C/C/C combination added up to almost 88% of the genotypes across the breeding stages while the A/T/T represented a little over 10%. The remaining 6 off-combinations summed up to less than 2% of the interactions. Although the allele frequencies were consistent with expectations under an epistatic model, the fact that interaction 2D–3B included only a single marker on chromosome segment 3B also led us to reject this potential three-way hypothesis as well.

## Discussion

### Random Genetic Drift Alone Does Not Create Strong and Consistent Interchromosomal LD

Strong linkage disequilibrium (LD) between pairs of markers—particularly when physically distant—can arise through random genetic drift, an effect that is more pronounced in small populations (Charlesworth et al., 2003; Charlesworth, 2009; Kliman et al., 2008; Rothammer et al., 2013). In this study, comparisons between theoretical and empirical R^2^ distributions showed that families simulated under the sole influence of drift consistently exhibited lower maximum interchromosomal LD values than empirical families, both when considering pairwise interactions present in at least two families (Figure 2) and when pooling all families for analysis (Figure 3). Importantly, even under conditions that favored the chance appearance of strong LD — smaller simulated family size (n = 15) and a fourfold increase in the total number of simulated families— the simulations produced no pairwise interactions that met the thresholds used to identify candidate interactions empirically. These findings indicate that random genetic drift alone is unlikely to create strong and consistent interchromosomal LD patterns across independent families.

### Strong interchromosomal LD could be created from selection of epistatic interactions

The poor performance of elite lines as parents is hypothesized to be related to epistatic effects. Although estimating epistatic variance is difficult because of non-orthogonality with additive effects (Tessele et al., 2024; Raffo et al., 2022; Vitezica et al., 2017), selection signatures of epistasis are expected to exist in breeding programs. In this study, 25 breeding families were utilized to identify 19 regions in strong interchromosomal LD which are hypothesized to be selection signatures of epistasis. Interchromosomal LD patterns were also observed in tetraploid wheat (Laido et al 2014) and common beans (Rossi et al., 2009; Diniz et al., 2018), but no inferences regarding the genetic mechanism leading to interchromosomal LD were drawn.

The allele frequency distribution of candidate interactions resembled that expected under the influence of sign epistasis, which falsifies the additive hypothesis. Similar results were also observed in wild barley, where the cross of plants based on spatial distance resulted in co-adapted gene complexes fixed in locally abundant genotypes (Volis et al., 2011). In two diverse populations of barley, several generations of natural selection resulted in two out of the 16 potential combinations of gametes (4^2^) as the most frequent, which presented perfectly complementary gametic types (Clegg et al. 1972), suggesting natural selection acted on coadapted (Dobzhansky, 1970) multilocus units.

As an attempt of disentangling epistasis from drift, an independent set of 15 validation families was utilized to further test the consistency of the candidate interactions, which falsified seven interactions. As the strength of genetic drift effects is expected to be especially predominant in smaller populations (Charlesworth et al., 2003; Charlesworth, 2009; Kliman et al., 2008; Rothammer et al. 2013), the assembly of allelic combinations resembling selection for sign epistasis in the discovery stage could still have been a consequence of genetic drift. Even though our simulation results suggested otherwise, random genetic drift has been attributed to the emergence of apparent selection signatures between pairs of SNPs in simulation studies (Id-Lahoucine et al., 2019; Vilas et al., 2012).

To mitigate the impact of genetic drift, a significantly larger dataset spanning a decade of breeding records was analyzed, leading to the rejection of four interactions. Among the remaining eight interactions, 2A-2D, 2D-3B, 2D-4A, and 3B-4A had a single allelic combination representing close to 90% of the total allelic combinations, contradicting the assumption of comparable adaptability between opposing allelic combinations (Wight, 1932). However, this observed phenomenon may relate to founder effects (James, 1971), where a restriction in allele frequency within the founder germplasm results in the spread of one of allelic combination in the breeding program. In that sense, the balanced frequency of opposed allelic combinations in interactions 1A-4A, 6B-6D, 3B-3D, and 2B-5B within an elite breeding program is surprising, considering that recycling lines and extensive utilization of good parental lines is expected to favor one epistatic combination over generations. For this reason, it is expected that the number and extent of sign epistatic interactions should be much larger when considering different breeding programs, as a function of founder effects, aka founder germplasm.

Even though the allele frequencies and interchromosomal linkage blocks observed for interactions 2D–3B, 3B–4A, 2B–5B, 3B–3D and 6B-6Da were consistent with expectations under the epistasis hypothesis, these interactions were not considered reliable for the purposes of this study. The polyploid nature of wheat and its large genome (∼17 Gb) enriched in repetitive DNA (Alipour et al., 2019) increase the likelihood of alignment errors during the GBS pipeline. Because each of these candidate interactions involved at least one genomic segment represented by a single marker, it is possible that the apparent strong linkage arose from misalignment rather than true biological signal. By contrast, we found no evidence to falsify interactions 1A–4A, 2D–4A, and 2D–3B, which therefore remain strong candidates for epistatic selection.

### Selection Against Unfavorable Epistatic Combinations and Its Implications for Wheat Breeding

The near-complete recovery of opposed allelic combinations as observed for interaction 1A-4A suggests artificial selection alone may not be the primary driver of distorted allele frequencies. Instead, it is possible that natural selection is responsible for recovering favorable allelic combinations, as observed in barley and wild barley populations (Volis et al., 2011; Clegg et al. 1972), through a nearly lethal response to off-combinations, similar to hybrid necrosis in wheat (Tsunewaki, 1960) or through embryo lethality, as reported in wheat-rye hybrids (Tikhenko et al., 2005; Tikhenko et al., 2010; Tikhenko et al., 2017). Anecdotal evidence on this hypothesis has been observed in single seed descent families in the Kansas State wheat breeding program. In this speed breeding pipeline, the quantity of seed recovered after three generations of inbreeding is often significantly lower than expected based solely on poor levels of germination. This could be an indication that a varying numbers of epistatic interactions, as interactions 1A-4A, 6B-6D, and 3B-3D, segregate in the families and plants homozygotes to off-combinations may experience a severe setback on performance which eventually leads to premature death or reproductive incapability. Populations with a higher number of segregating epistatic interactions, if additionally segregating to interactions 2B-5B, 2A-2D, and 2D-3B, for instance, may consequently end up with fewer viable plants.

A practical parallel can be drawn with the well-known *Rht-1* genes in wheat, which confer reduced responsiveness to gibberellic acid and lead to a semi-dwarf phenotype (Gale and Marshall, 1973; Pinthus et al., 1989). Although most modern cultivars carry either *Rht-B1b* or *Rht-D1b* (Evans, 1998), the combination of both alleles results in a dwarf phenotype that is highly undesirable. Similarly, when breeding programs combine genotypes that achieve a target phenotype through distinct genetic mechanisms, unfavorable allelic interactions can emerge, effectively reducing the viable population size and constraining breeding progress, and reduce the rate of genetic gain in the long term (Fall et al., 2025).

One alternative to overcome this challenge is to constrain the genetic diversity of the breeding program with the fixation of one of the interacting alleles, converting epistatic variance into additive variance (Hill et al., 2008; Technow et al., 2021). In the candidate interaction 1A-4A, fixing the A allele in chromosome 1A causes the allele in chromosome 4A to behave additively, where the T allele is always positive, and C is always negative (Figure 5 A, C, and E). This ensures that the selection pressure is constant for the alleles in the interacting locus and increases the probability of having one favorable combination fixed.

### Sign Epistasis Creates a Rugged Surface on the Fitness Landscape

The possible existence of di-genic sign epistasis, as observed in interaction 1A-4A (Figure 5A), implies that transitioning from one superior combination to another requires two allelic substitutions, passing through a transitional state of inferior performance (Wright, 1988; Wade, 2002). The fitness landscape of the potential tri-genic sign epistasis interaction reported in this study has a valley between the two optimum allelic combinations (Figure 6B). And, if extended to higher order epistasis and more gene interactions (Figure 6C), it indicates that different combinations of alleles can present similar levels of adaptation to the same environmental conditions (Wright, 1988). The entire field of allelic combinations and respective levels of adaptation can be greatly simplified through the fitness landscape metaphor (Figure 6D), as proposed in the shifting balance theory (Wright, 1932), where hills and valleys represent high and low levels of adaptation, respectively. The fitness landscape, as a metaphor, should not be taken literally (Wade, 2002), as it is inadequate to represent higher order epistatic interactions (Wright, 1932). It does, however, provides a powerful simplified representation of complex genetic systems and how populations navigate the rugged surface during evolution. The concept of evolution under the shifting balance theory has striking similarities with the structure of breeding programs (Technow et a., 2021) and could be leveraged to better understand epistasis in the face of managing breeding germplasm pools.

### Breeding Strategies on a Rugged Fitness Landscape

A rugged surface on the fitness landscape has implications for breeding programs as the strategy used to manage the germplasm pool can lead to higher or lower levels of epistatic variance in breeding families. The operation of a closed breeding program inevitably reduces the genetic diversity in the germplasm pool and, under the influence of drift and selection, leads to the fixation of alleles involved in sign epistatic interactions. This process converts epistatic variance into additive variance and increases the potential of adaptation of the breeding program (populations) to local environments (Wade and Goodnight, 1998), as less complex genetic systems have higher heritability and respond better to selection. This can be visualized in Figure 6D as the *black squares* or *blue circles* genotypes, located in independent hills and assumed to comprise separate breeding programs. Alternatively, operating a breeding program with a large genetic base may reintroduce epistatic variance when recycling lines from within the breeding program, which can be depicted as the *white star* genotypes in Figure 6D. The cross of parental lines that belong to different adaptive peaks leads to a phenomenon called F_2_ breakdown (Whitlock et al., 1995), where the mean performance of the population is lower than the midparent value. This is a consequence of disrupting, in the offspring, the contrasting epistatic interactions that were homozygous in each parental line. Breeding families with high levels of epistatic variance present a low correlation between early and late generations (Upadhyaya and Nigam, 1998; Humphrey et al., 1969), indicating a limited response to early generation selection. If late generation selection is adopted, the quick pace of inbreeding and the commonly small population size of breeding families causes both favorable and unfavorable combinations to be fixed, reducing the probability of assembling optimal allelic combinations.

One common feature of breeding programs is the exchange of germplasm, where elite genotypes are anticipated to transmit superior allelic combinations from one breeding pool to another (Wright, 1932; Technow et al., 2021). The success of introducing germplasm on a rugged fitness landscape scenario depends on having both parental lines on the same adaptive peak, that is, sharing the same founding allelic combinations, otherwise, F_2_ breakdown may also be observed in breeding families. For navigating on multiple adaptive peaks, broad genetic base programs may be more suitable for introducing external germplasm as there is more than one set of founding allelic combinations that could match the germplasm to be introduced. Although normally used as a strategy to introduce a few alleles in the background of a recurrent line (Hospital, 2005), backcrossing can be leveraged by closed breeding programs to recover the epistatic framework of the recurrent line. Backcrossing constraints epistatic variance, increasing additive variance and the response to selection in breeding populations. For a comprehensive discussion of the implications of sign epistasis from the perspective of the fitness landscape metaphor, view Tessele et al. (2024) and Holland (2001).

## Conclusion

This study investigated whether selection signatures consistent with sign epistasis exist within wheat breeding families, aiming to expand the understanding of how epistasis may underlie the occasional poor performance of elite lines as parents. A genome-wide analysis detected 19 candidate interchromosomal interactions with abnormally strong linkage disequilibrium whose allelic patterns were consistent with sign epistasis and refuted the competing additive hypothesis. To avoid circular inference and minimize bias from genetic drift, two independent validation sets were analyzed. Eleven interactions were falsified, and five were excluded because they involved a single marker on one of the interacting chromosomes, where strong LD and allele frequency distortions may have arisen from genotyping-by-sequencing (GBS) alignment artifacts. There was not sufficient evidence to reject the sign epistasis hypothesis for three candidate interactions with the current dataset and analysis. The possibility existence of sign epistasis could be responsible for a reduction in the effective population size in crosses with contrasting sign epistasis combinations, and for the eventual failure to recycle elite lines in the breeding program as epistatic variance is reestablished in the offspring. The potential presence of a rugged fitness landscape underscores the importance of adequately managing epistatic variance within breeding programs for efficiently developing elite germplasm.

## Data Availability

Marker information of discovery and validation populations in HapMap format is included in files ‘Discovery-Populations.hmp.txt’ and ‘Validation-Populations.hmp.txt’. The code to run the PCA analysis to eliminate outliers within families is found in ‘PCA.R’. The code used to calculate linkage disequilibrium is found in ‘C-Linkage_Disequilibrium_Loops.bach’. The code used to calculate the 2D-LD plots in discovery and validation families and the control plot is found in the `*_2D_LDplot.R’ files. The ‘Circleze.R’ file has the code used to plot interchromosomal LD with chromosomes displayed circularly. The file ‘Simulation.R’ contains the R code used to simulate families under the sole influence of drift. Supplemental material is available at Figshare. The raw genotypic data, available at Figshare, will be deposited at NCBI if/upon the manuscript is accepted for publication.

