## Supplemental Tables for "Interchromosomal Linkage Disequilibrium Analysis Reveals Strong Indications of Sign Epistasis in Wheat Breeding Families"

**Table 1.** Summary of the main SNPs in strong interchromosomal linkage disequilibrium.

| Chr 1 | SNP 1 | Chr 2* | SNP 2 | Allelic Combinations** | Families Segregating | # Families fixed | | # Pairwise Interactions | # Unique Markers*** | Outcome of hypothesis |
|---|---|---|---|---|---|---|---|---|---|---|
| | | | | | | Comb 1 | Comb 2 | | | |
| **1A** | 565519207 | **4A** | 598236311 | A/T - C/C | 15 | 8 | 2 | 39 | 19 – 5 | Not falsified |
| **2D** | 91221807 | **4A** | 693794455 | C/C - A/T | 12 | 13 | 0 | 146 | 10 – 33 | Not falsified |
| **2A** | 713142609 | **2D** | 572257428 | A/C - G/G | 12 | 1 | 12 | 11 | 4 – 7 | Not falsified |
| **2D** | 91221807 | **3B** | 713887174 | C/C - A/T | 10 | 15 | 0 | 6 | 6 – 1 | Possibility of GBS alignment error |
| **3B** | 713887174 | **4A** | 693794455 | C/C - T/T | 10 | 15 | 0 | 18 | 1 – 18 | Possibility of GBS alignment error |
| **2B** | 25458554 | **5B** | 600423012 | A/A - G/G | 15 | 9 | 1 | 6 | 1 – 6 | Possibility of GBS alignment error |
| **3B** | 752546905 | **3D** | 567747632 | C/T - T/A | 12 | 3 | 10 | 1 | 1 – 1 | Possibility of GBS alignment error |
| **6B** | 26204619 | **6Da** | 14436093 | C/A - T/G | 17 | 1 | 7 | 1 | 1 – 1 | Possibility of GBS alignment error |
| **1B** | 659660316 | **6B** | 669333749 | A/T - G/C | 4 | 20 | 1 | 6 | 2 – 3 | Potential drift in breeding program |
| **5D** | 378314409 | **7B** | 661308545 | T/A - A/G | 4 | 21 | 0 | 2 | 1 – 2 | Potential drift in breeding program |
| **3A** | 17107524 | **3D** | 9881568 | A/G - T/A | 4 | 21 | 0 | 2 | 2 – 1 | Potential drift in breeding program |
| **1A** | 589041708 | **1B** | 683324328 | T/A - C/C | 12 | 13 | 0 | 6 | 1 – 6 | Potential drift in breeding program |
| **5A** | 598520962 | **6B** | 81973027 | C/A - G/G | 8 | 1 | 16 | 5 | 3 – 2 | Potential drift in breeding program |
| **6A** | 611814110 | **7A** | 4327054 | T/G - CC | 6 | 19 | 0 | 11 | 1 – 11 | Potential drift in breeding program |
| **5A** | 687959438 | **7A** | 17199897 | T/C - C/T | 12 | 13 | 0 | 3 | 1 – 3 | Allele frequency in validation panel |
| **1BL** | 146796395 | **1D** | 391069881 | T/A - G/C | 12 | 13 | 0 | 1 | 1 – 1 | Allele frequency in validation panel |
| **1BS** | 39577562 | **1D** | 391069881 | A/A - T/C | 12 | 13 | 0 | 1 | 1 – 1 | Allele frequency in validation panel |
| **2B** | 12856952 | **7A** | 704484934 | T/C - C/T | 5 | 20 | 0 | 2 | 2 – 1 | Allele frequency in validation panel |
| **6B** | 33136603 | **6Db** | 19567942 | C/G - A/T | 4 | 21 | 0 | 2 | 1 – 2 | Allele frequency in validation panel |

**Two interactions involved the chromosome 6BS and the same region on chromosome 6D. To distinguish between interactions, the letters a and b was added to each of them.*

*** The allelic combinations represents the two opposing allelic combinations under selection. For example, in the first interaction (1A-4A), SNP S1A_565519207 had the alleles A and C, while SNP S4A_598236311 had the alleles T and C. The four allelic combinations were: A/T, A/C, C/T, and C/C.*

**** The number on the left represents the number of unique markers found in Chr 1 of that specific interaction, while the number on the right represents the number of unique markers found in Chr 2.*

**A**

| | Family 1 | | Family 2 | | Family n | | | Sum Families | |
|---|---|---|---|---|---|---|---|---|---|
| | $B_1$ | $B_2$ | $B_1$ | $B_2$ | $B_1$ | $B_2$ | | $B_1$ | $B_2$ |
| $A_1$ | 48 | 2 | 63 | 1 | 21 | 1 | → | 132 | 4 |
| $A_2$ | 1 | 49 | 1 | 35 | 0 | 78 | | 2 | 162 |
| | $R^2$ =0.89 | | $R^2$ =0.92 | | $R^2$ =0.94 | | | $R^2$ =0.92 | |

**B**

| | | | | | | | | | |
|---|---|---|---|---|---|---|---|---|---|
| $A_1$ | 48 | 2 | 1 | 35 | 0 | 78 | → | 49 | 115 |
| $A_2$ | 1 | 48 | 63 | 1 | 21 | 1 | | 85 | 50 |
| | $R^2$ =0.89 | | $R^2$ =0.92 | | $R^2$ =0.94 | | | $R^2$ =0.11 | |

**C**

| | | | | | | | | | |
|---|---|---|---|---|---|---|---|---|---|
| $A_1$ | 100 | 0 | 100 | 0 | 0 | 0 | → | 200 | 0 |
| $A_2$ | 0 | 0 | 0 | 0 | 0 | 100 | | 0 | 100 |
| | | | | | | | | $R^2$ =1 | |

**Table 2. Illustration of how allele frequencies in pairwise alleles lead to strong interchromosomal LD within and across families.** (A) Consistent pairing of $A_1B_1$ and $A_2B_2$ alleles within families, as expected under the influence of epistasis, results in strong linkage when families are analyzed individually or when allele frequencies from all families are combined. (B) Random genetic drift can cause families to share the same pairwise interactions in strong linkage but with different preferential allelic combinations, leading to weak linkage when the families are analyzed together. (C) Population structure can produce strong linkage between pairs of alleles when families are analyzed collectively, while within-family linkage disequilibrium is absent because the interactions are fixed.

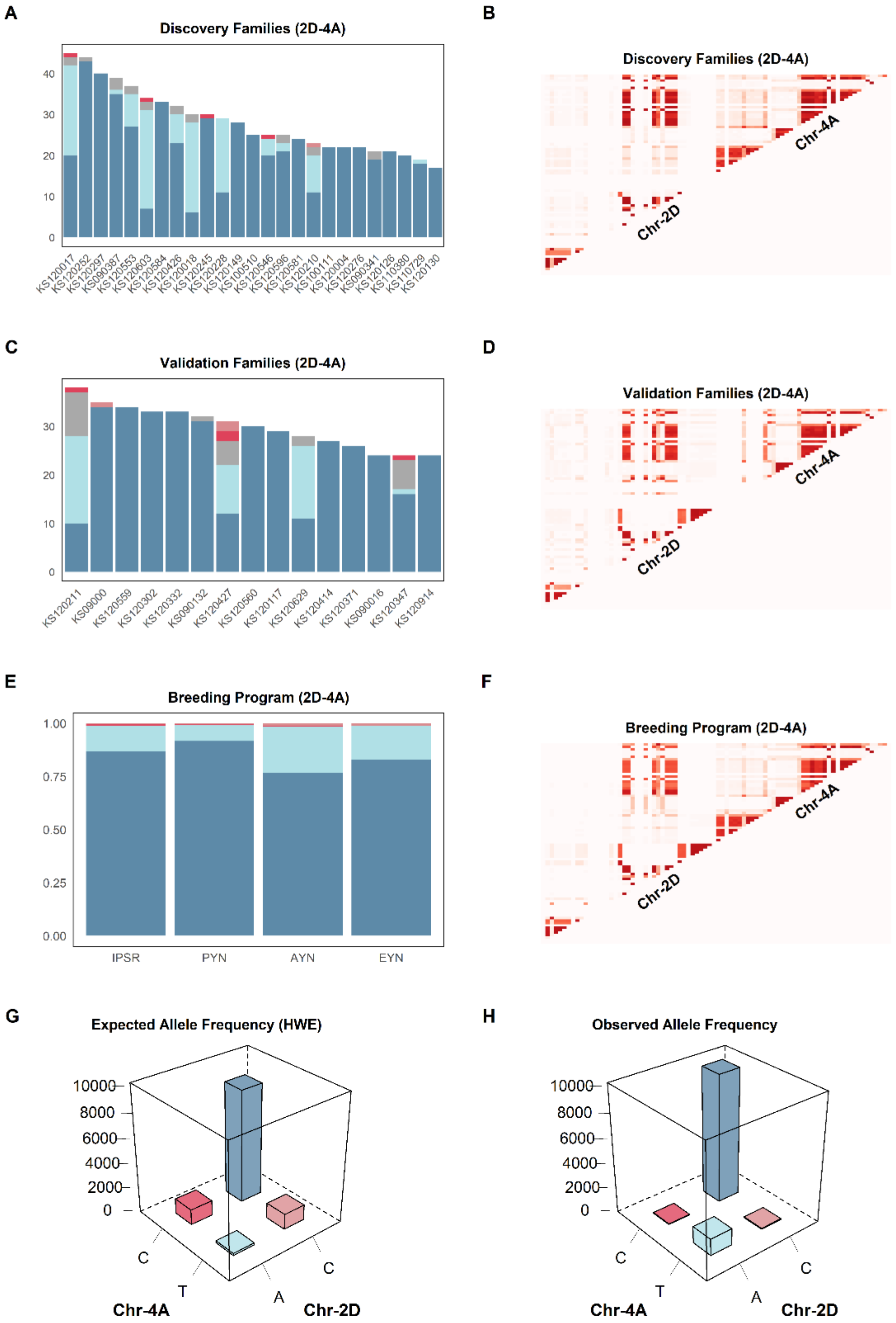


**Figure 1** Testing independent breeding panels for sign epistasis. Allele frequency distribution (left column) and heatmaps of 2D LD plots (right column) of the candidate epistatic interaction involving segments of chromosomes **2D** and **4A**. **A**, **C** and **E** display the allele frequency distribution from the discovery and validation panels, and from the last four stages of the breeding pipeline, respectively. **B**, **D** and **F** shows the 2D LD plot from discovery and validation panels, and from the last four stages of the breeding pipeline, respectively. **G** displays the expected allele frequency distribution in the entire breeding program under Hardy-Weinberg Equilibrium and **H** shows the observed allele frequency distribution.

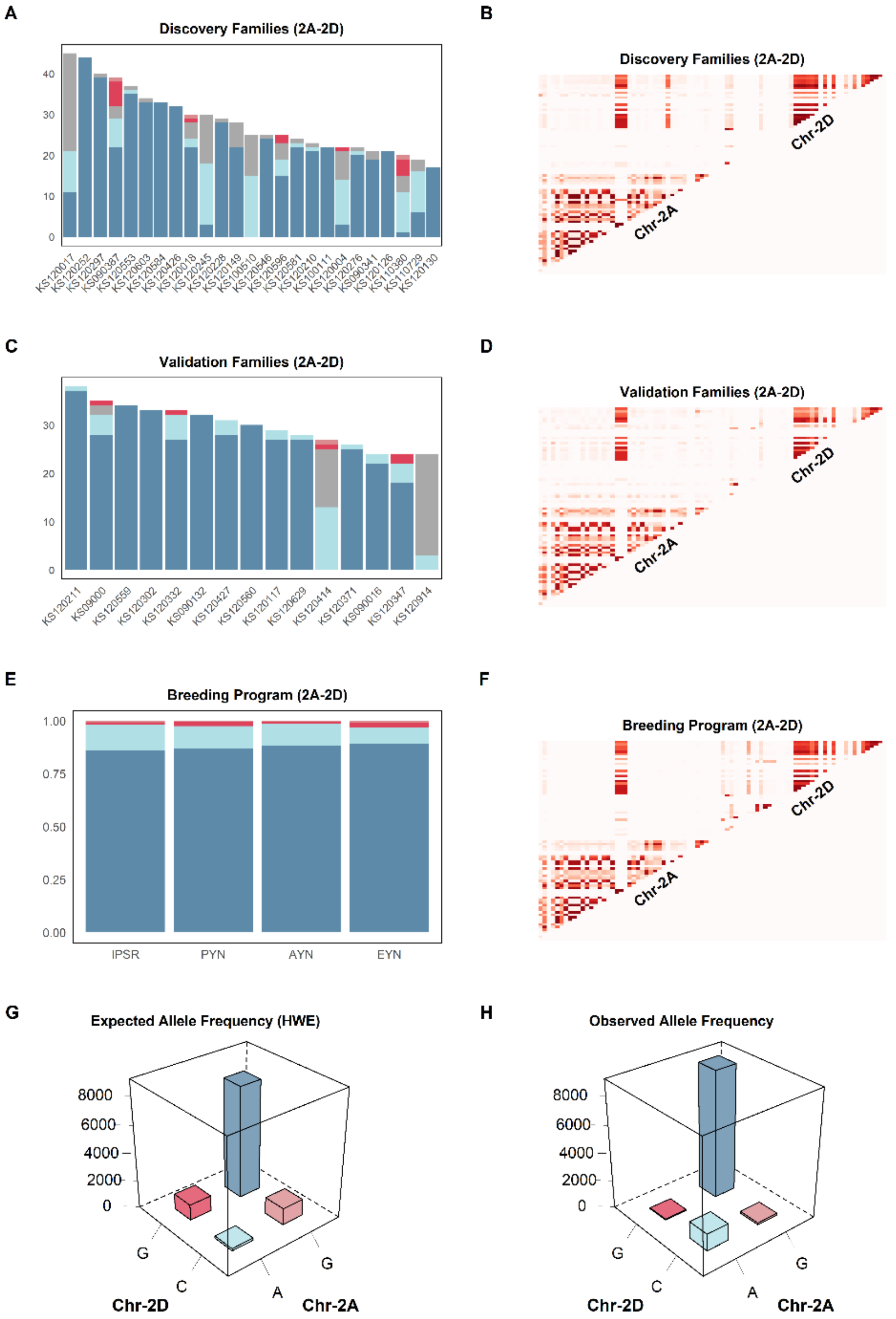


**Figure 2** Testing independent breeding panels for sign epistasis. Allele frequency distribution (left column) and heatmaps of 2D LD plots (right column) of the candidate epistatic interaction involving segments of chromosomes **2A** and **2D**. **A**, **C** and **E** display the allele frequency distribution from the discovery and validation panels, and from the last four stages of the breeding pipeline, respectively. **B**, **D** and **F** shows the 2D LD plot from discovery and validation panels, and from the last four stages of the breeding pipeline, respectively. **G** displays the expected allele frequency distribution in the entire breeding program under Hardy-Weinberg Equilibrium and **H** shows the observed allele frequency distribution.

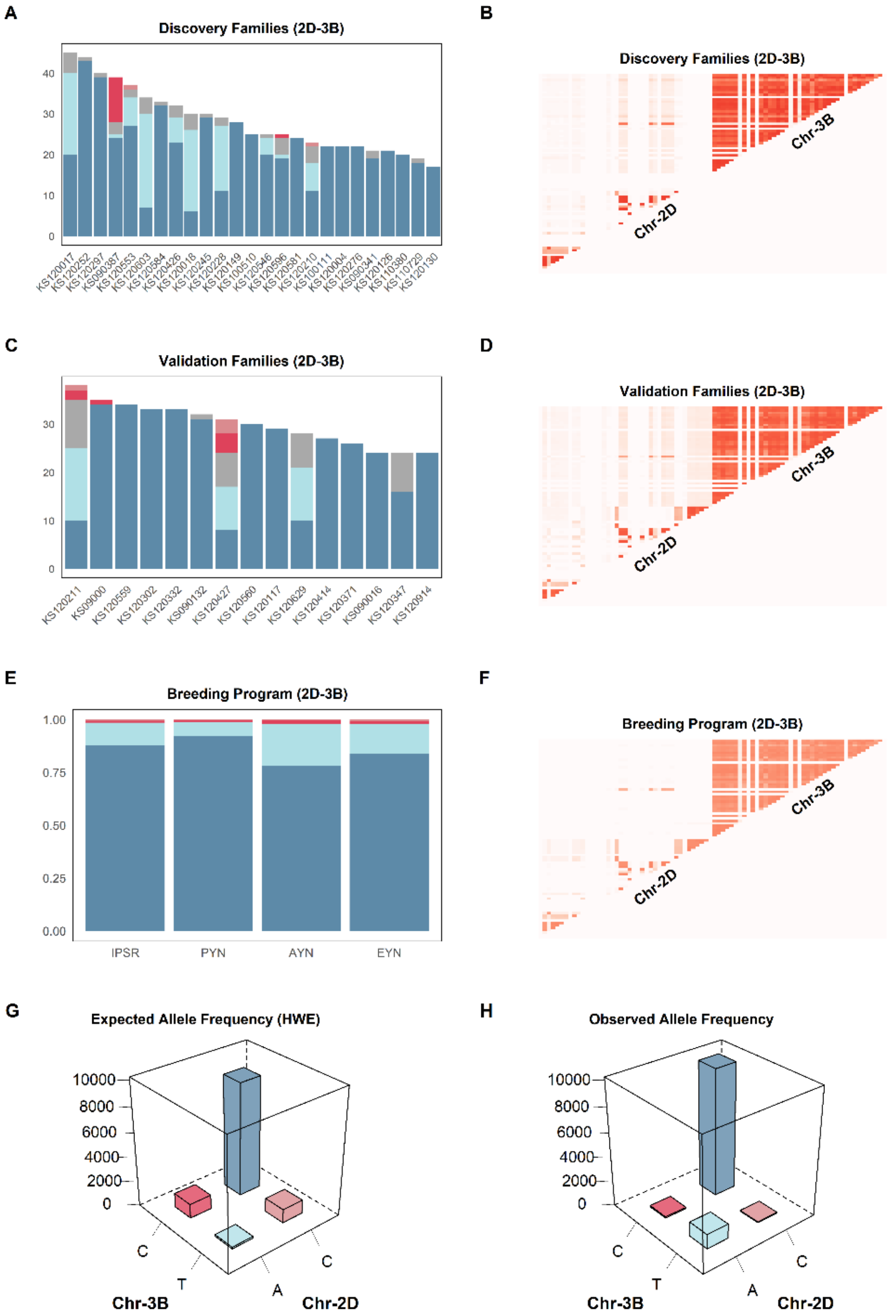


**Figure 3** Testing independent breeding panels for sign epistasis. Allele frequency distribution (left column) and heatmaps of 2D LD plots (right column) of the candidate epistatic interaction involving segments of chromosomes **2D** and **3B**. **A**, **C** and **E** display the allele frequency distribution from the discovery and validation panels, and from the last four stages of the breeding pipeline, respectively. **B**, **D** and **F** shows the 2D LD plot from discovery and validation panels, and from the last four stages of the breeding pipeline, respectively. **G** displays the expected allele frequency distribution in the entire breeding program under Hardy-Weinberg Equilibrium and **H** shows the observed allele frequency distribution.

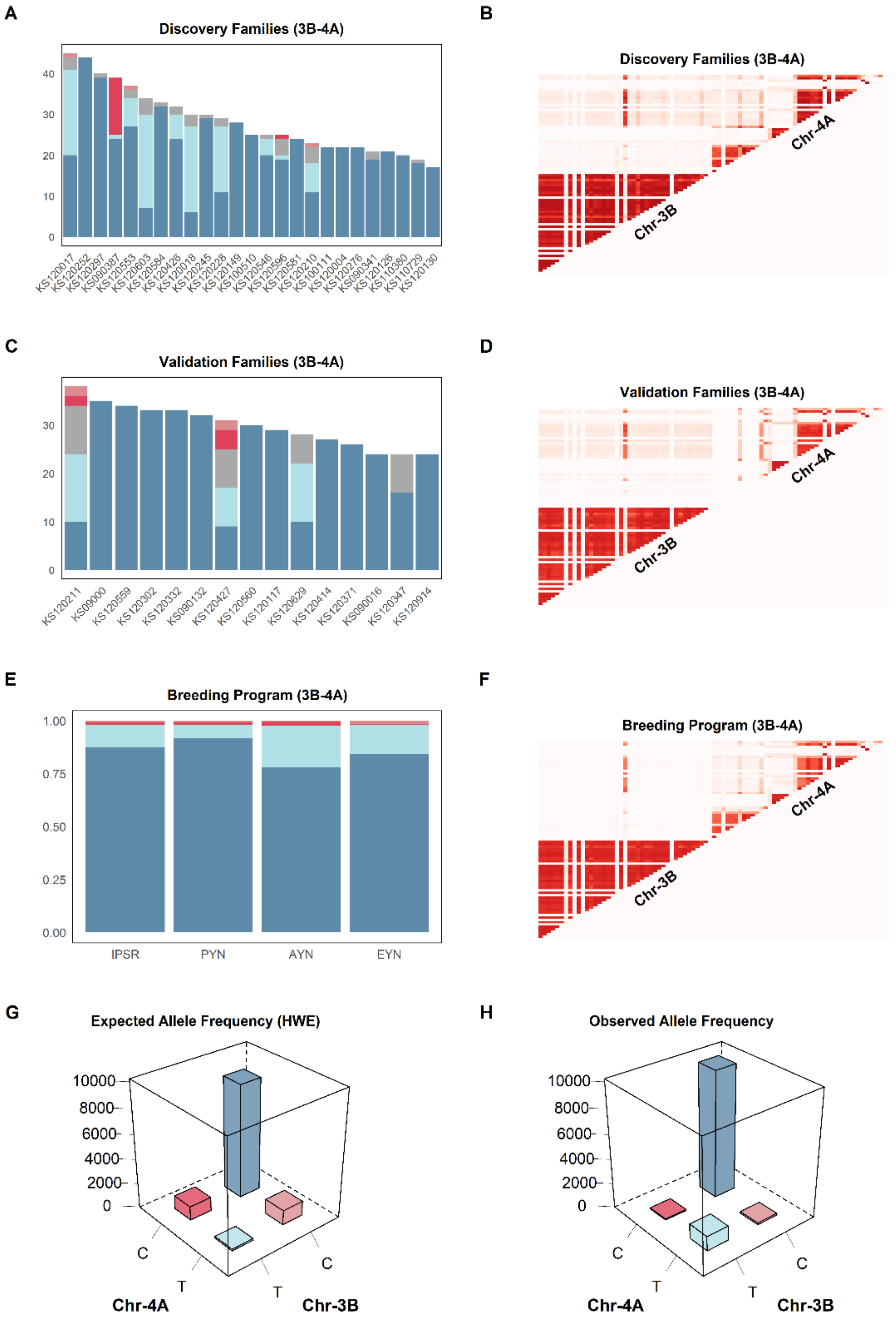


**Figure 4** Testing independent breeding panels for sign epistasis. Allele frequency distribution (left column) and heatmaps of 2D LD plots (right column) of the candidate epistatic interaction involving segments of chromosomes **3B** and **4A**. **A**, **C** and **E** display the allele frequency distribution from the discovery and validation panels, and from the last four stages of the breeding pipeline, respectively. **B**, **D** and **F** shows the 2D LD plot from discovery and validation panels, and from the last four stages of the breeding pipeline, respectively. **G** displays the expected allele frequency distribution in the entire breeding program under Hardy-Weinberg Equilibrium and **H** shows the observed allele frequency distribution.

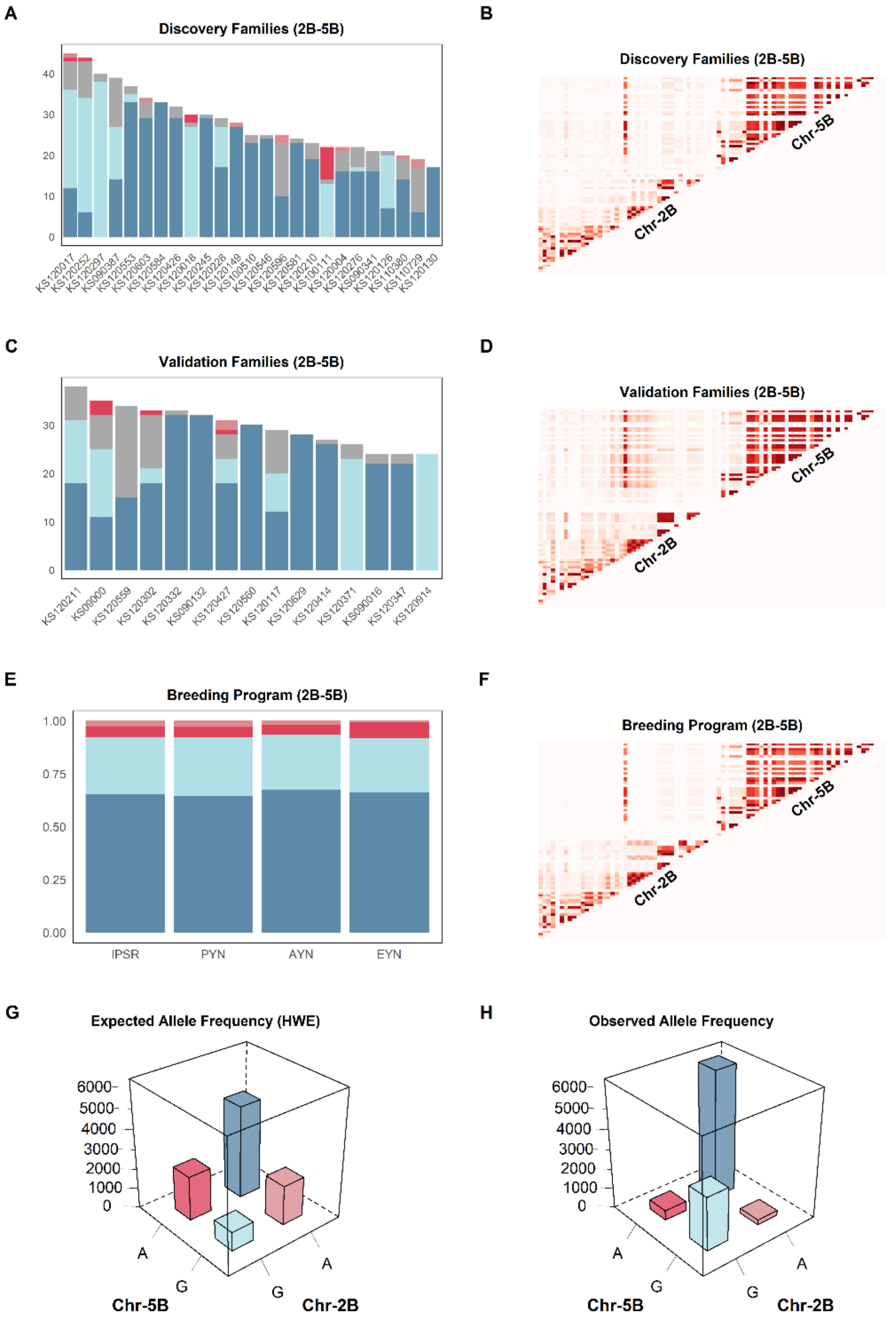


**Figure 5** Testing independent breeding panels for sign epistasis. Allele frequency distribution (left column) and heatmaps of 2D LD plots (right column) of the candidate epistatic interaction involving segments of chromosomes **2B** and **5B**. **A**, **C** and **E** display the allele frequency distribution from the discovery and validation panels, and from the last four stages of the breeding pipeline, respectively. **B**, **D** and **F** shows the 2D LD plot from discovery and validation panels, and from the last four stages of the breeding pipeline, respectively. **G** displays the expected allele frequency distribution in the entire breeding program under Hardy-Weinberg Equilibrium and **H** shows the observed allele frequency distribution.

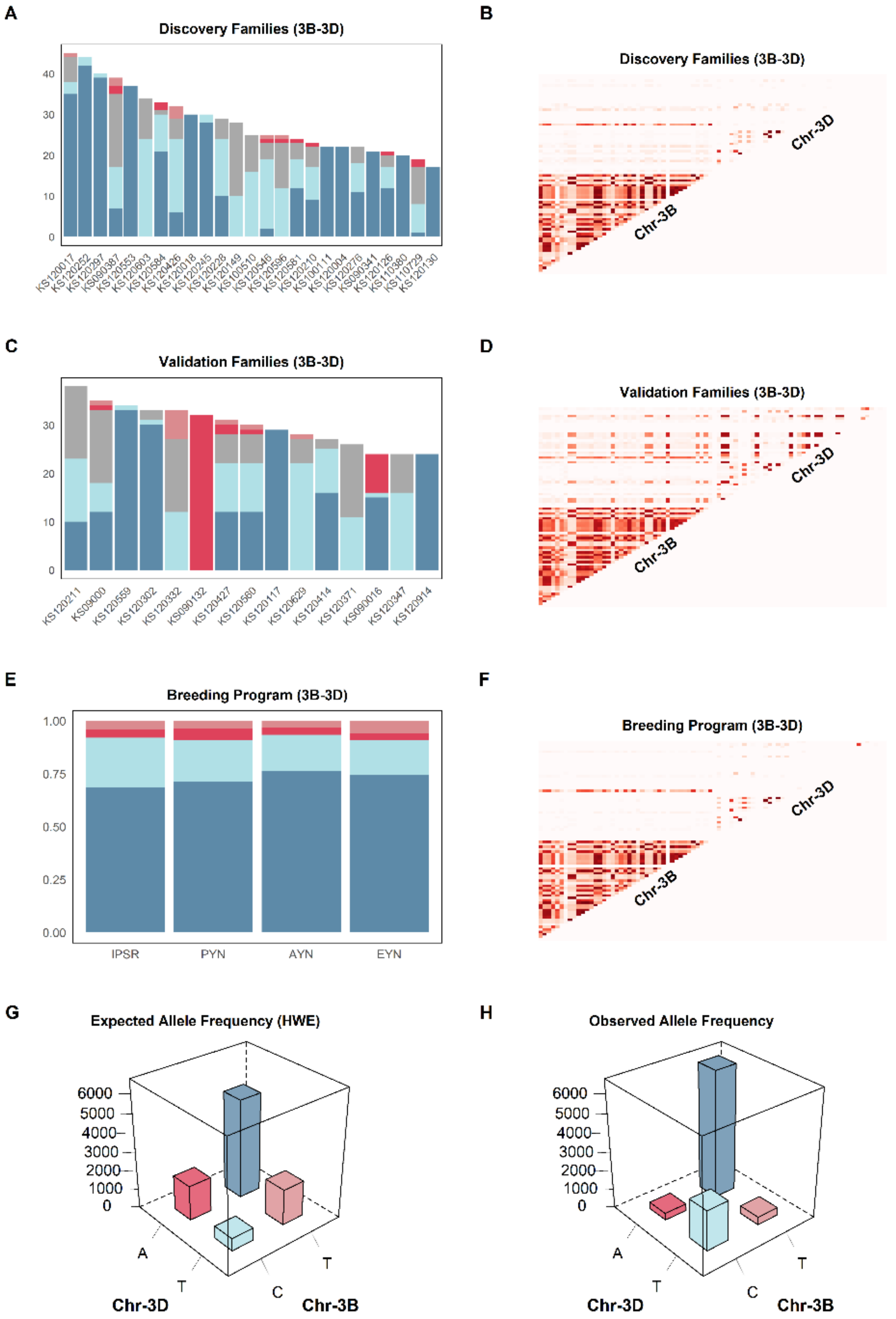


**Figure 6** Testing independent breeding panels for sign epistasis. Allele frequency distribution (left column) and heatmaps of 2D LD plots (right column) of the candidate epistatic interaction involving segments of chromosomes **3B** and **3D**. **A**, **C** and **E** display the allele frequency distribution from the discovery and validation panels, and from the last four stages of the breeding pipeline, respectively. **B**, **D** and **F** shows the 2D LD plot from discovery and validation panels, and from the last four stages of the breeding pipeline, respectively. **G** displays the expected allele frequency distribution in the entire breeding program under Hardy-Weinberg Equilibrium and **H** shows the observed allele frequency distribution.

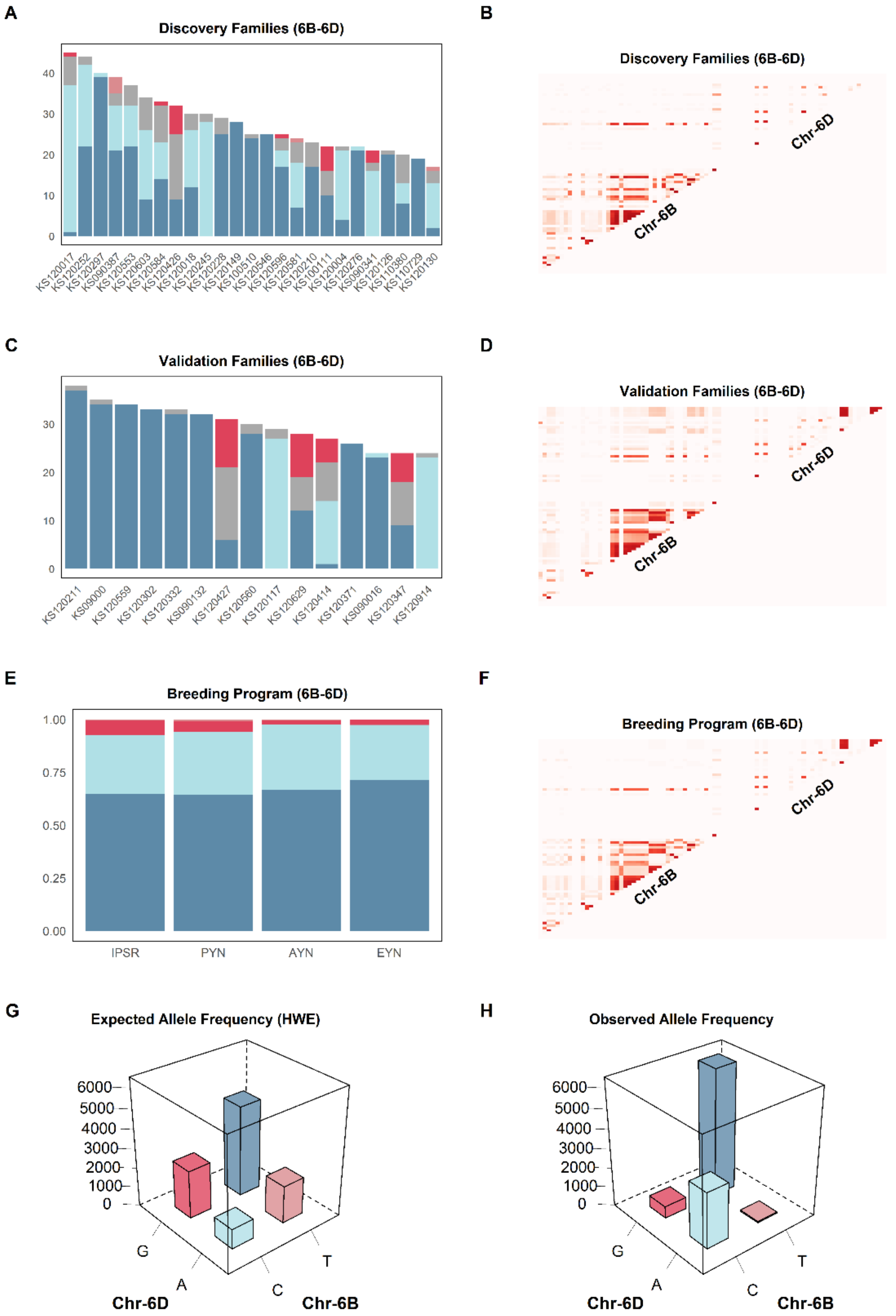


**Figure 7** Testing independent breeding panels for sign epistasis. Allele frequency distribution (left column) and heatmaps of 2D LD plots (right column) of the candidate epistatic interaction involving segments of chromosomes 6B and 6Da. A, C and E display the allele frequency distribution from the discovery and validation panels, and from the last four stages of the breeding pipeline, respectively. B, D and F shows the 2D LD plot from discovery and validation panels, and from the last four stages of the breeding pipeline, respectively. G displays the expected allele frequency distribution in the entire breeding program under Hardy-Weinberg Equilibrium and H shows the observed allele frequency distribution.

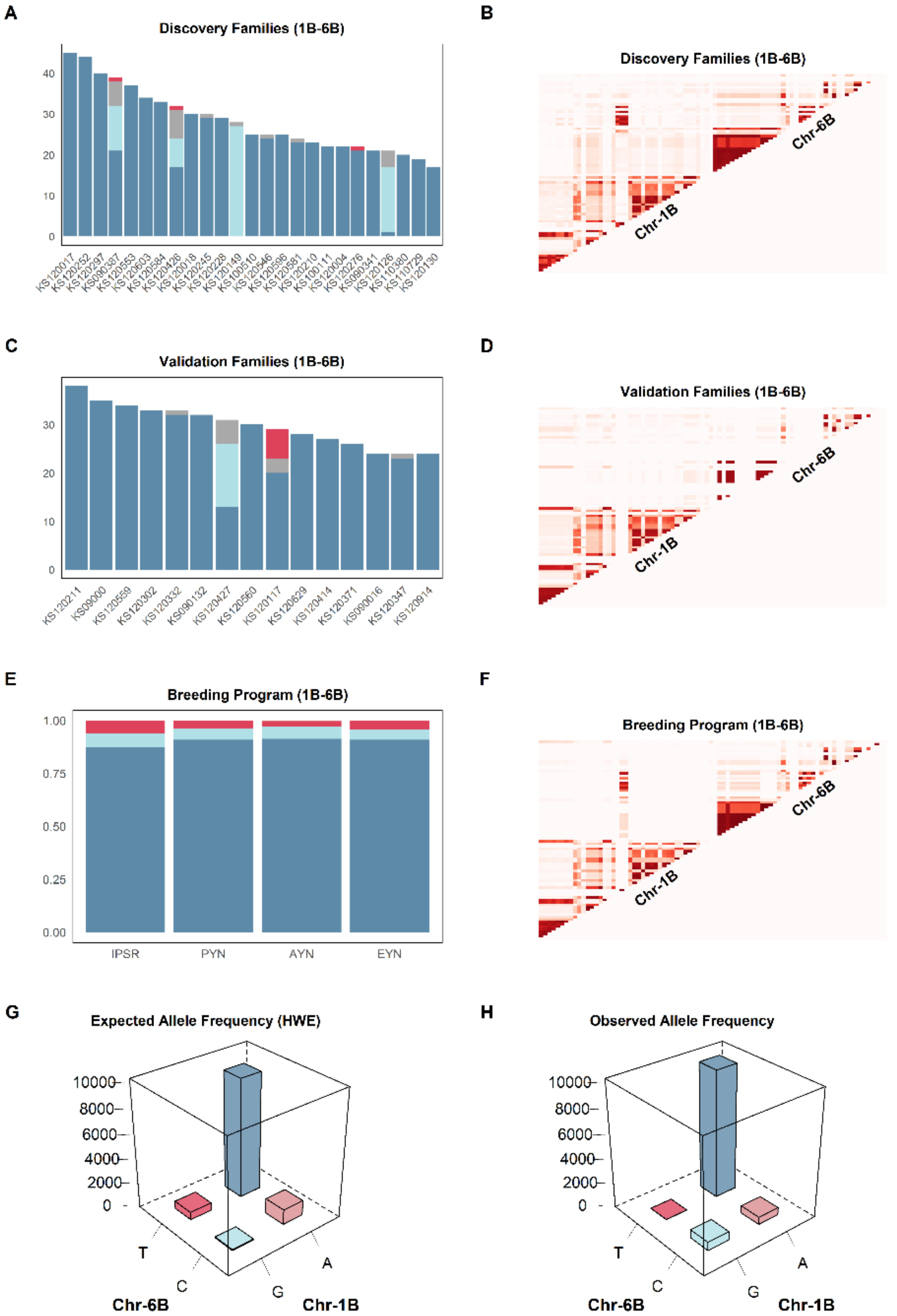


**Figure 8** Testing independent breeding panels for sign epistasis. Allele frequency distribution (left column) and heatmaps of 2D LD plots (right column) of the candidate epistatic interaction involving segments of chromosomes 1B and 6B. A, C and E display the allele frequency distribution from the discovery and validation panels, and from the last four stages of the breeding pipeline, respectively. B, D and F shows the 2D LD plot from discovery and validation panels, and from the last four stages of the breeding pipeline, respectively. G displays the expected allele frequency distribution in the entire breeding program under Hardy-Weinberg Equilibrium and H shows the observed allele frequency distribution.

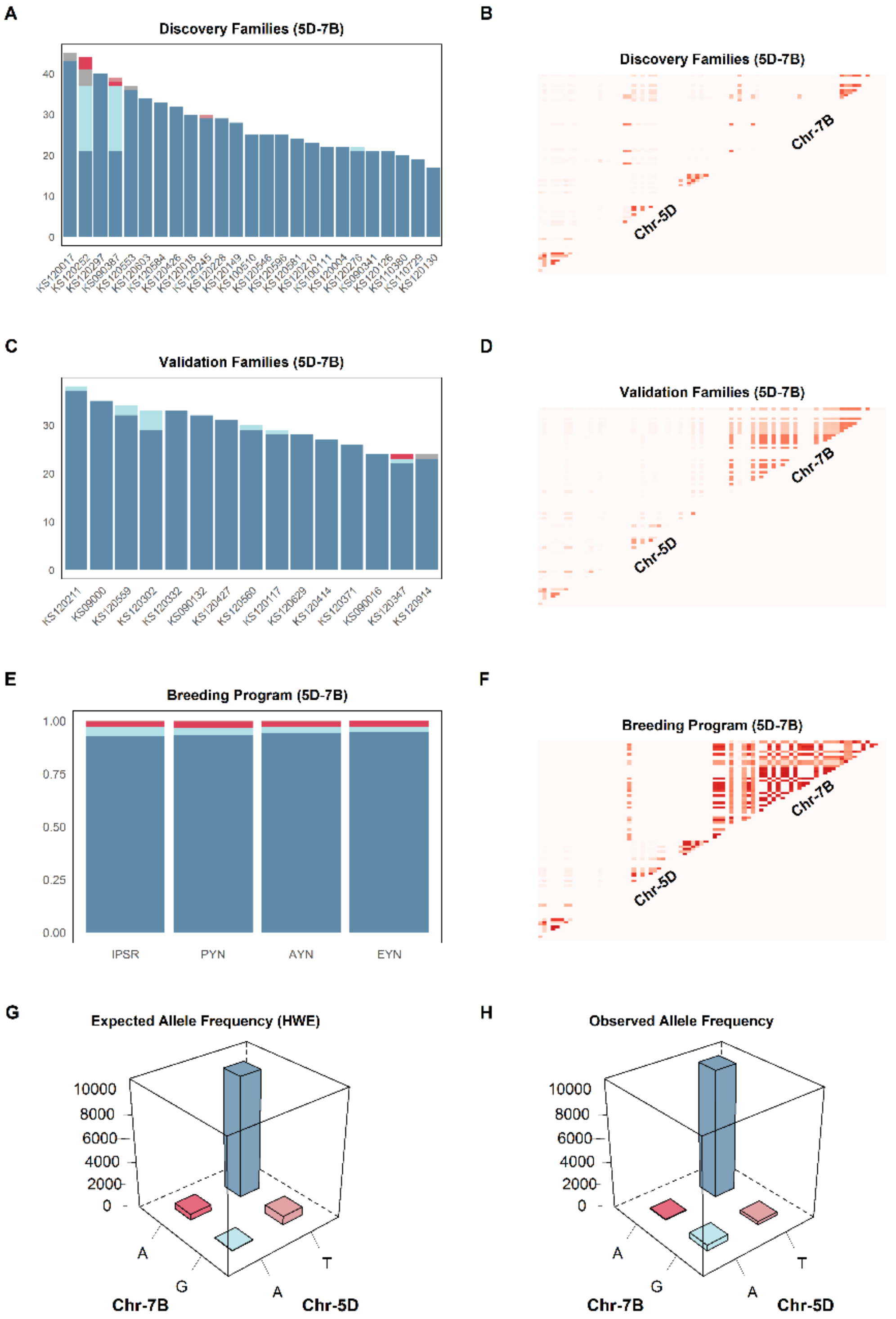


**Figure 9** Testing independent breeding panels for sign epistasis. Allele frequency distribution (left column) and heatmaps of 2D LD plots (right column) of the candidate epistatic interaction involving segments of chromosomes **5D** and **7B**. **A**, **C** and **E** display the allele frequency distribution from the discovery and validation panels, and from the last four stages of the breeding pipeline, respectively. **B**, **D** and **F** shows the 2D LD plot from discovery and validation panels, and from the last four stages of the breeding pipeline, respectively. **G** displays the expected allele frequency distribution in the entire breeding program under Hardy-Weinberg Equilibrium and **H** shows the observed allele frequency distribution.

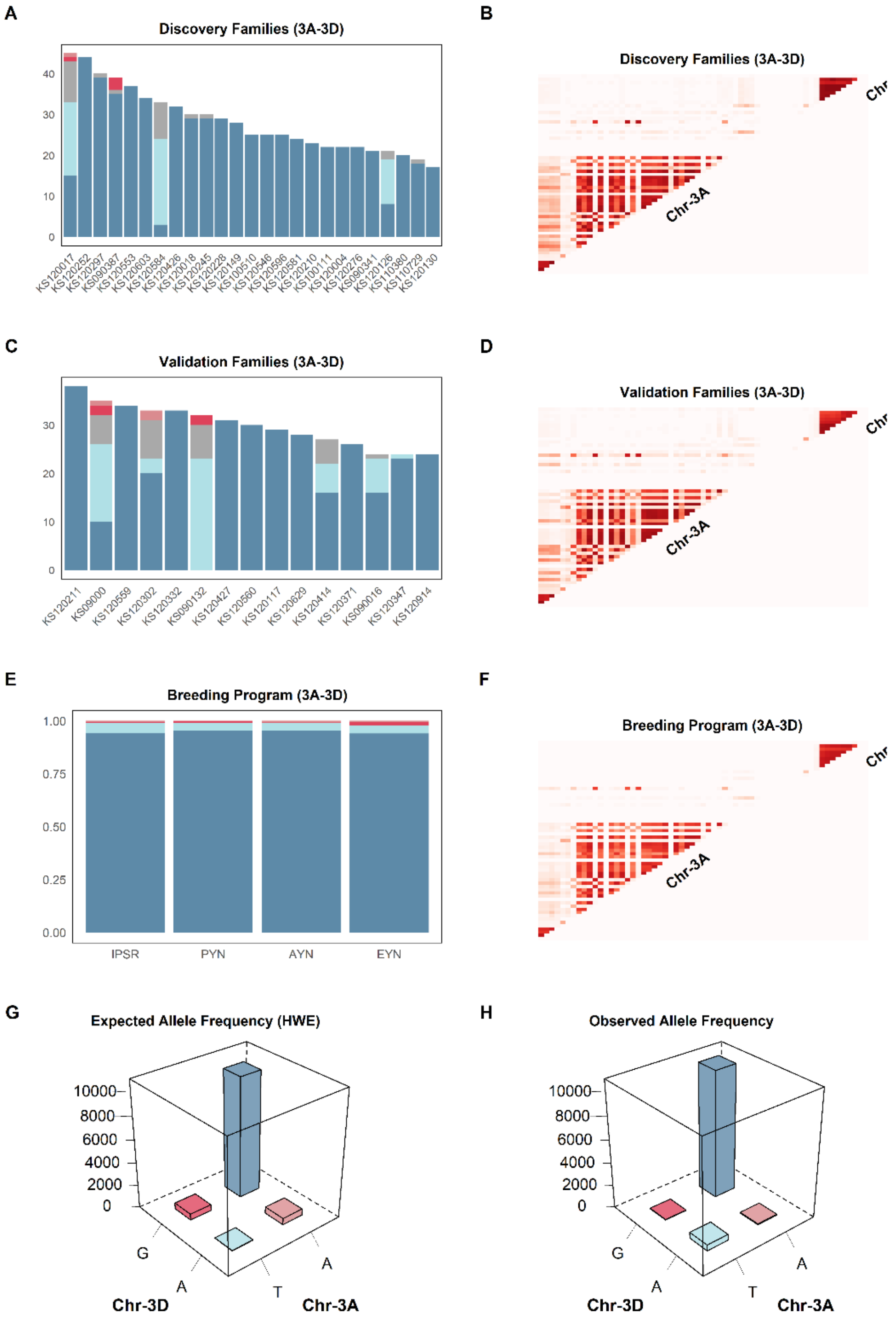


**Figure 10** Testing independent breeding panels for sign epistasis. Allele frequency distribution (left column) and heatmaps of 2D LD plots (right column) of the candidate epistatic interaction involving segments of chromosomes **3A** and **3D**. **A**, **C** and **E** display the allele frequency distribution from the discovery and validation panels, and from the last four stages of the breeding pipeline, respectively. **B**, **D** and **F** shows the 2D LD plot from discovery and validation panels, and from the last four stages of the breeding pipeline, respectively. **G** displays the expected allele frequency distribution in the entire breeding program under Hardy-Weinberg Equilibrium and **H** shows the observed allele frequency distribution.

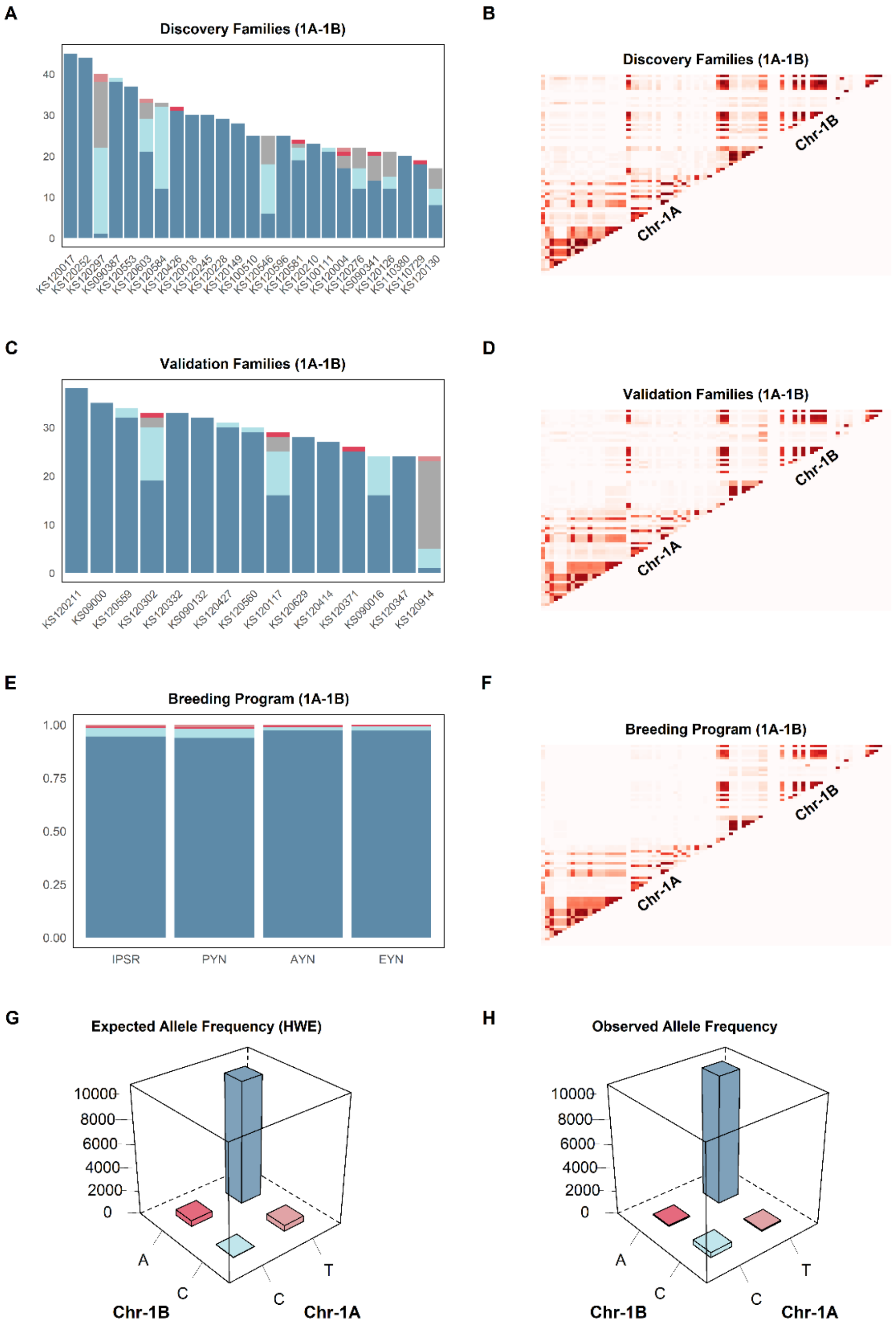


**Figure 11** Testing independent breeding panels for sign epistasis. Allele frequency distribution (left column) and heatmaps of 2D LD plots (right column) of the candidate epistatic interaction involving segments of chromosomes **1A** and **1B**. **A**, **C** and **E** display the allele frequency distribution from the discovery and validation panels, and from the last four stages of the breeding pipeline, respectively. **B**, **D** and **F** shows the 2D LD plot from discovery and validation panels, and from the last four stages of the breeding pipeline, respectively. **G** displays the expected allele frequency distribution in the entire breeding program under Hardy-Weinberg Equilibrium and **H** shows the observed allele frequency distribution.

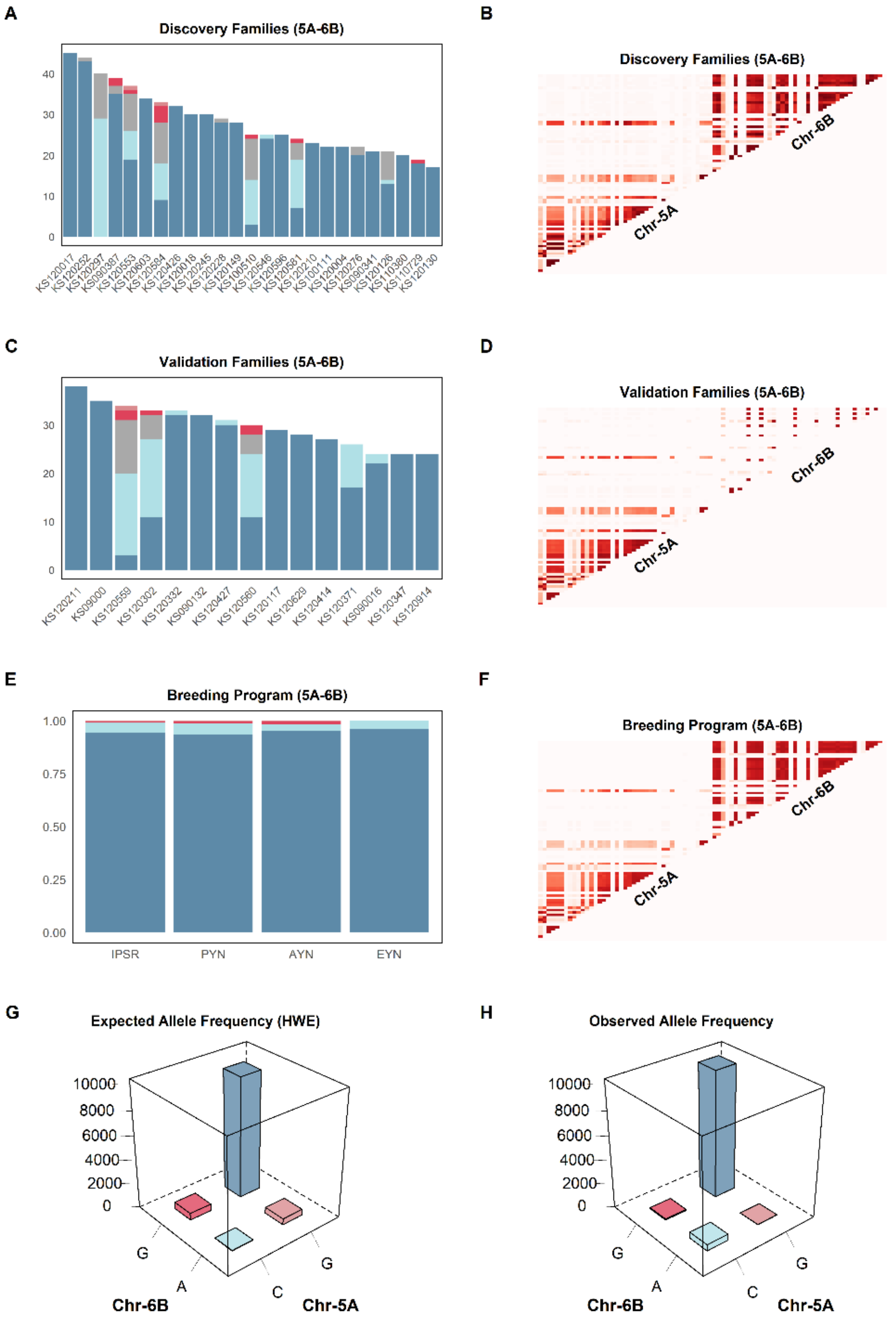


**Figure 12** Testing independent breeding panels for sign epistasis. Allele frequency distribution (left column) and heatmaps of 2D LD plots (right column) of the candidate epistatic interaction involving segments of chromosomes **5A** and **6B**. **A**, **C** and **E** display the allele frequency distribution from the discovery and validation panels, and from the last four stages of the breeding pipeline, respectively. **B**, **D** and **F** shows the 2D LD plot from discovery and validation panels, and from the last four stages of the breeding pipeline, respectively. **G** displays the expected allele frequency distribution in the entire breeding program under Hardy-Weinberg Equilibrium and **H** shows the observed allele frequency distribution.

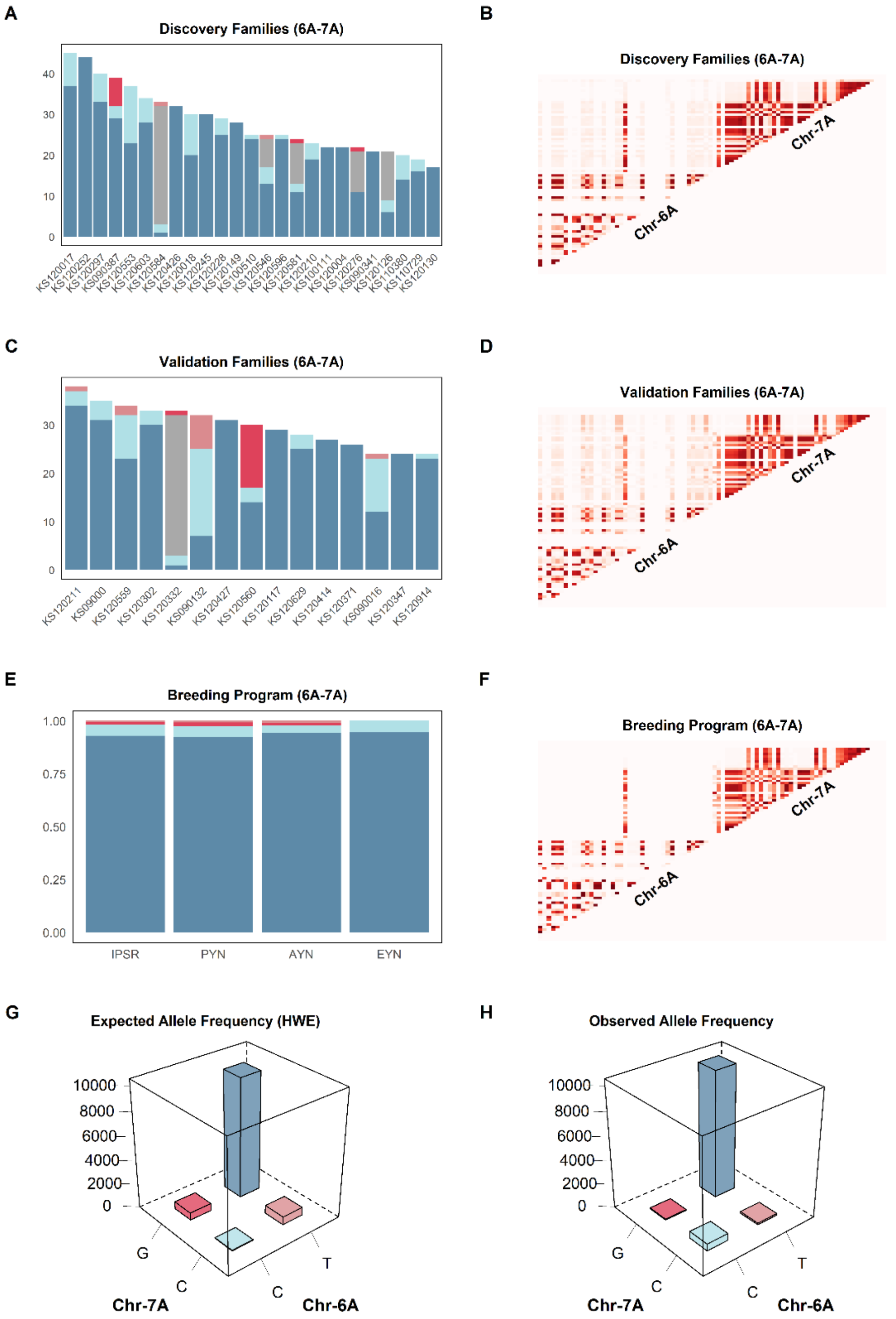


**Figure 13** Testing independent breeding panels for sign epistasis. Allele frequency distribution (left column) and heatmaps of 2D LD plots (right column) of the candidate epistatic interaction involving segments of chromosomes **6A** and **7A**. **A**, **C** and **E** display the allele frequency distribution from the discovery and validation panels, and from the last four stages of the breeding pipeline, respectively. **B**, **D** and **F** shows the 2D LD plot from discovery and validation panels, and from the last four stages of the breeding pipeline, respectively. **G** displays the expected allele frequency distribution in the entire breeding program under Hardy-Weinberg Equilibrium and **H** shows the observed allele frequency distribution.

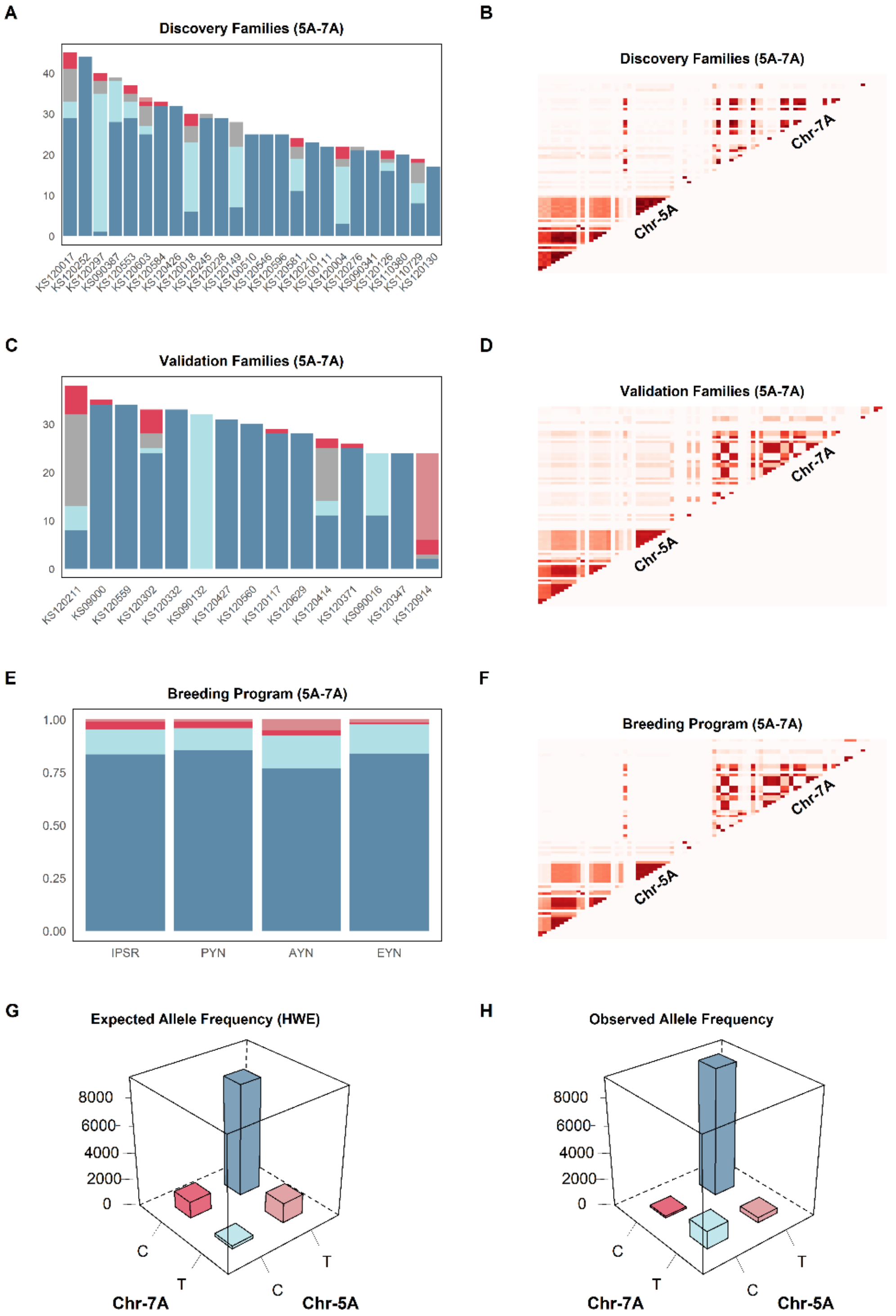


**Figure 14** Testing independent breeding panels for sign epistasis. Allele frequency distribution (left column) and heatmaps of 2D LD plots (right column) of the candidate epistatic interaction involving segments of chromosomes **5A** and **7A**. **A**, **C** and **E** display the allele frequency distribution from the discovery and validation panels, and from the last four stages of the breeding pipeline, respectively. **B**, **D** and **F** shows the 2D LD plot from discovery and validation panels, and from the last four stages of the breeding pipeline, respectively. **G** displays the expected allele frequency distribution in the entire breeding program under Hardy-Weinberg Equilibrium and **H** shows the observed allele frequency distribution.

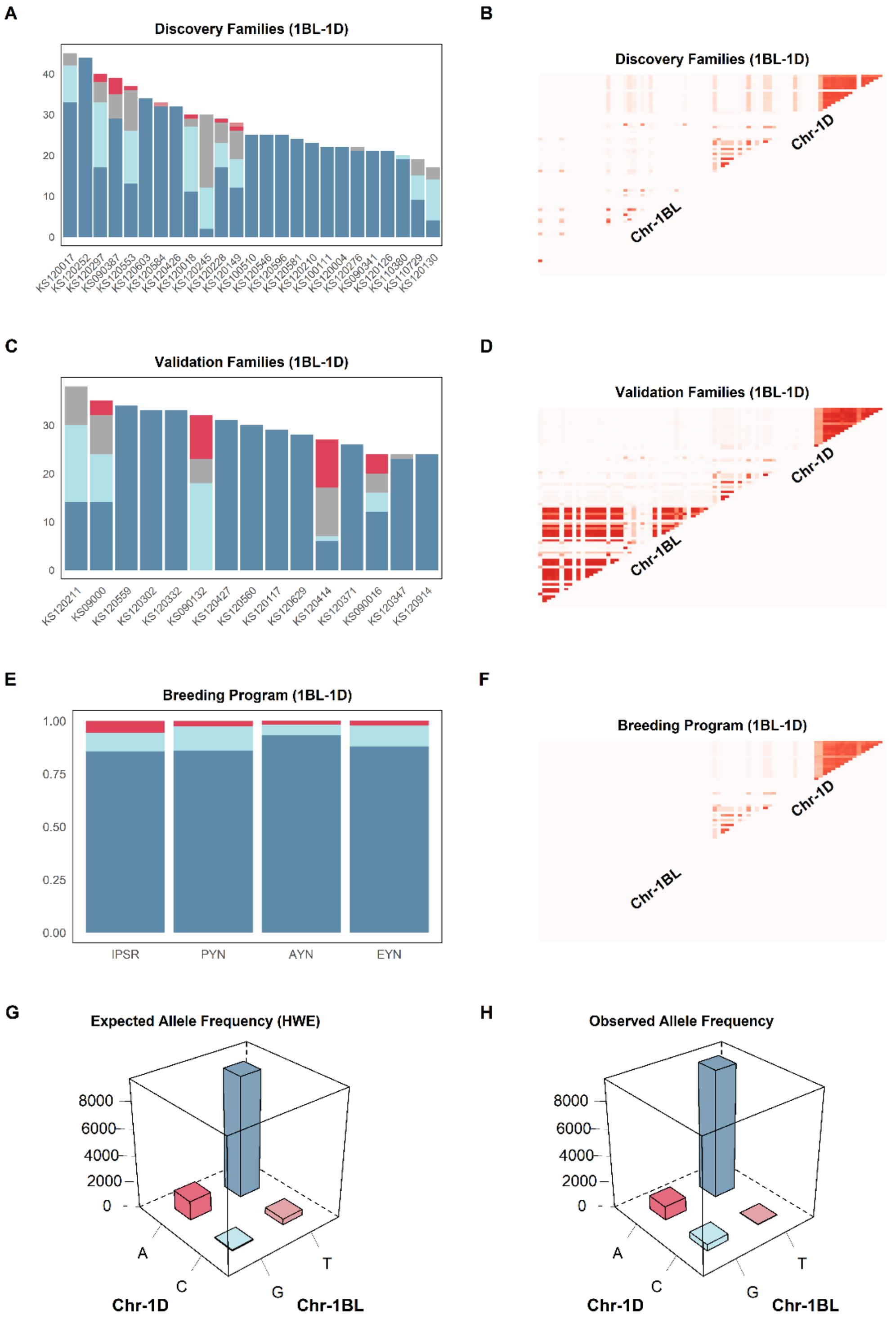


**Figure 15** Testing independent breeding panels for sign epistasis. Allele frequency distribution (left column) and heatmaps of 2D LD plots (right column) of the candidate epistatic interaction involving segments of chromosomes **1BL** and **1D**. **A**, **C** and **E** display the allele frequency distribution from the discovery and validation panels, and from the last four stages of the breeding pipeline, respectively. **B**, **D** and **F** shows the 2D LD plot from discovery and validation panels, and from the last four stages of the breeding pipeline, respectively. **G** displays the expected allele frequency distribution in the entire breeding program under Hardy-Weinberg Equilibrium and **H** shows the observed allele frequency distribution.

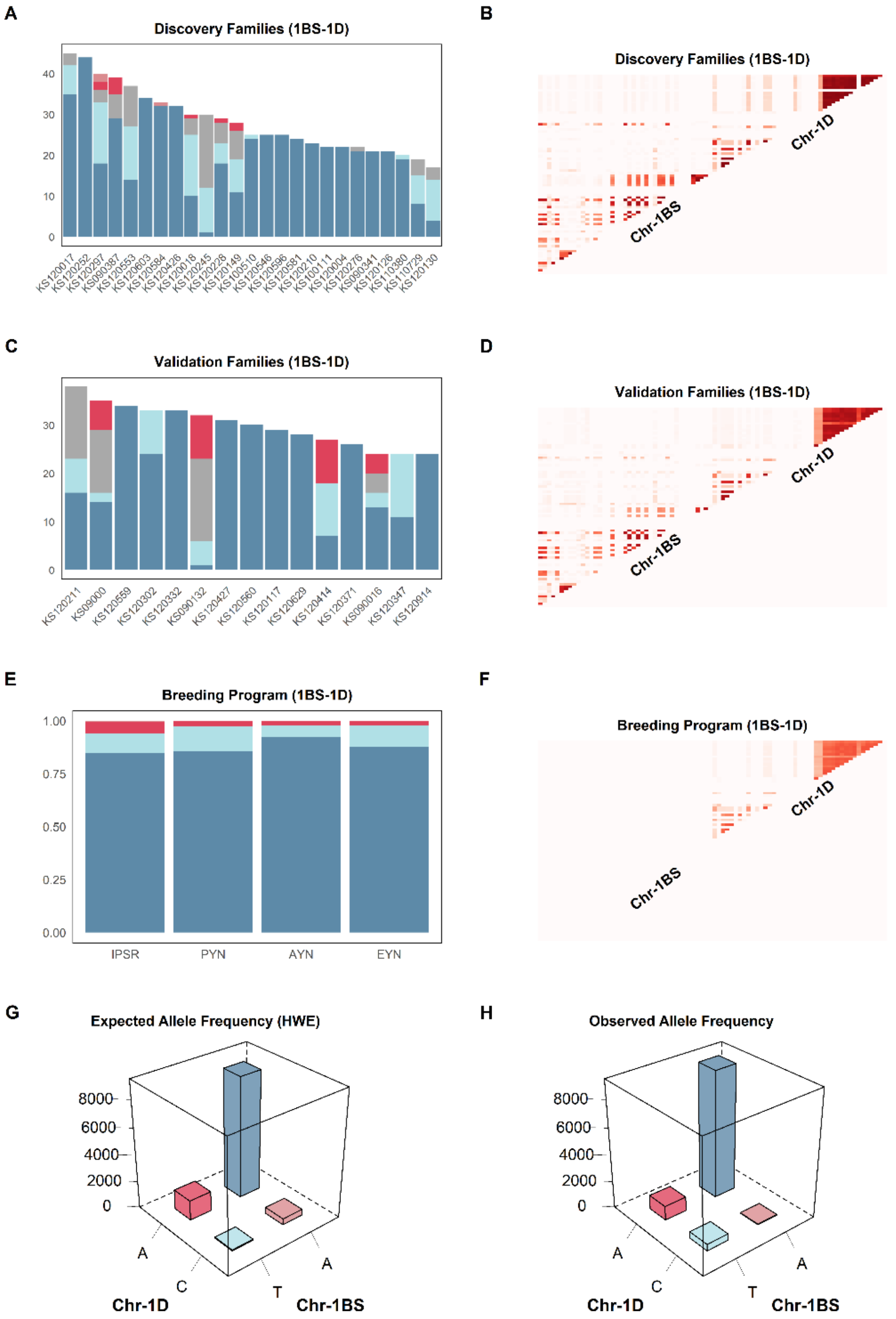


**Figure 16** Testing independent breeding panels for sign epistasis. Allele frequency distribution (left column) and heatmaps of 2D LD plots (right column) of the candidate epistatic interaction involving segments of chromosomes **1BS** and **1D**. **A**, **C** and **E** display the allele frequency distribution from the discovery and validation panels, and from the last four stages of the breeding pipeline, respectively. **B**, **D** and **F** shows the 2D LD plot from discovery and validation panels, and from the last four stages of the breeding pipeline, respectively. **G** displays the expected allele frequency distribution in the entire breeding program under Hardy-Weinberg Equilibrium and **H** shows the observed allele frequency distribution.

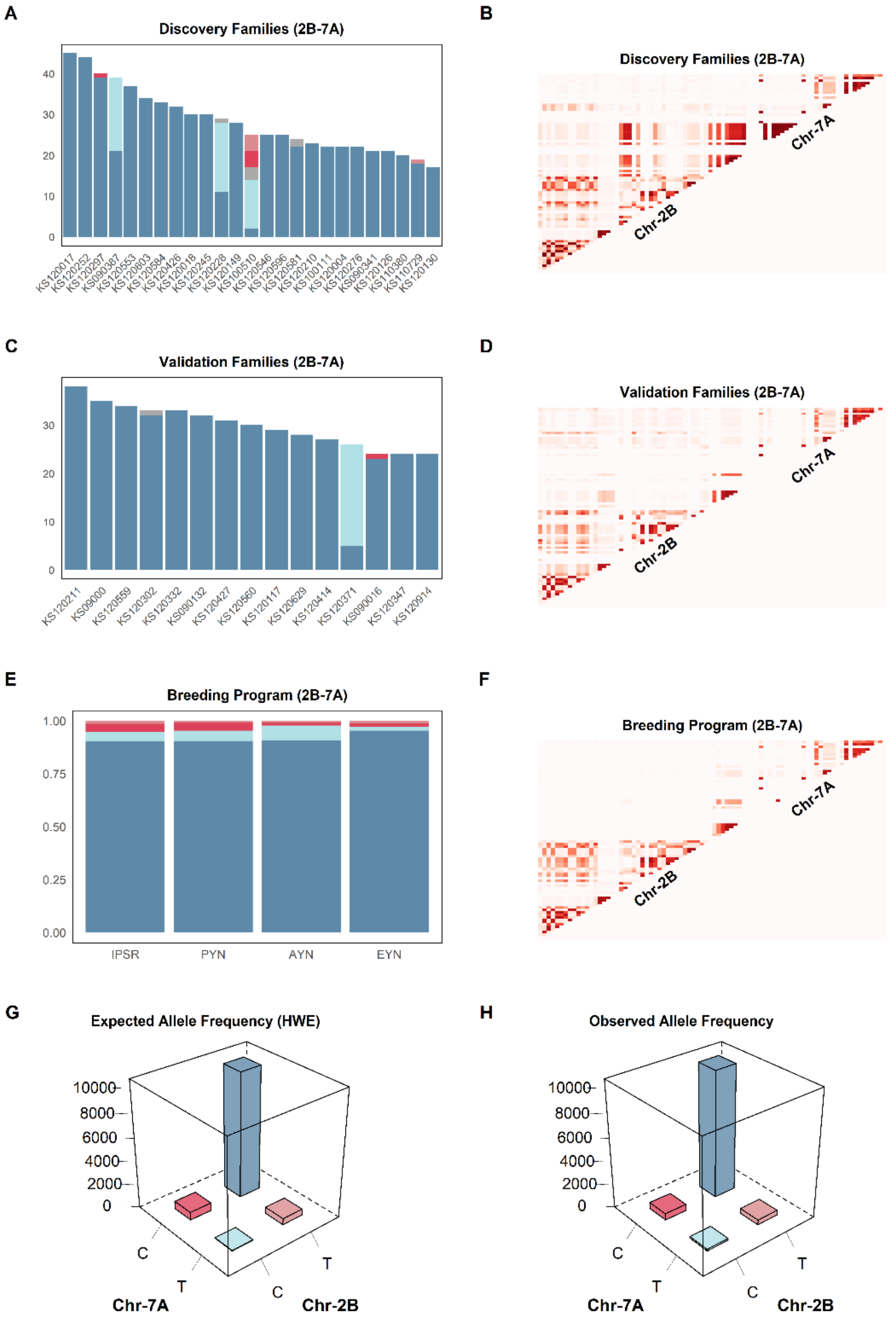


**Figure 17** Testing independent breeding panels for sign epistasis. Allele frequency distribution (left column) and heatmaps of 2D LD plots (right column) of the candidate epistatic interaction involving segments of chromosomes **2B** and **7A**. **A**, **C** and **E** display the allele frequency distribution from the discovery and validation panels, and from the last four stages of the breeding pipeline, respectively. **B**, **D** and **F** shows the 2D LD plot from discovery and validation panels, and from the last four stages of the breeding pipeline, respectively. **G** displays the expected allele frequency distribution in the entire breeding program under Hardy-Weinberg Equilibrium and **H** shows the observed allele frequency distribution.

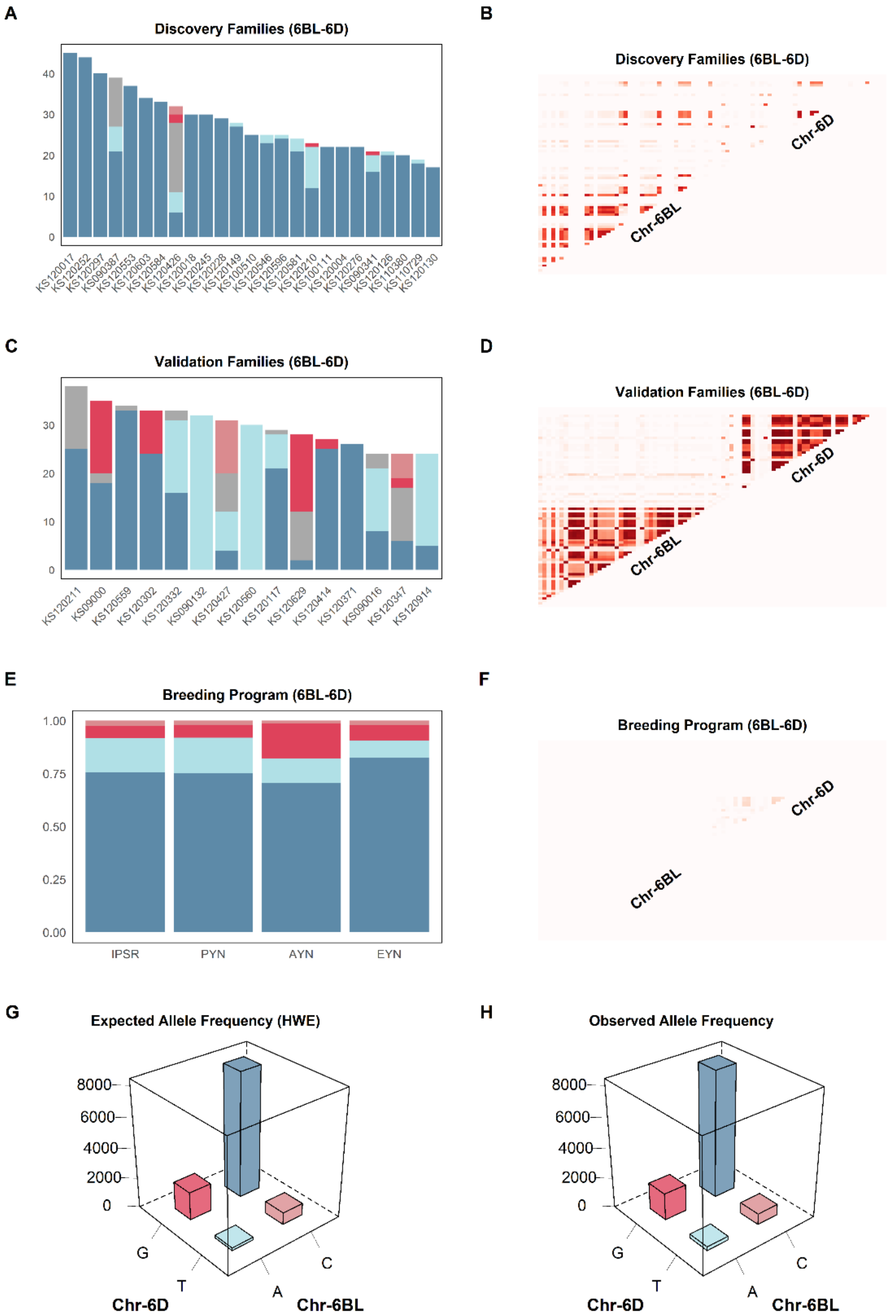


**Figure 18** Testing independent breeding panels for sign epistasis. Allele frequency distribution (left column) and heatmaps of 2D LD plots (right column) of the candidate epistatic interaction involving segments of chromosomes **6BL** and **6D**. **A**, **C** and **E** display the allele frequency distribution from the discovery and validation panels, and from the last four stages of the breeding pipeline, respectively. **B**, **D** and **F** shows the 2D LD plot from discovery and validation panels, and from the last four stages of the breeding pipeline, respectively. **G** displays the expected allele frequency distribution in the entire breeding program under Hardy-Weinberg Equilibrium and **H** shows the observed allele frequency distribution.

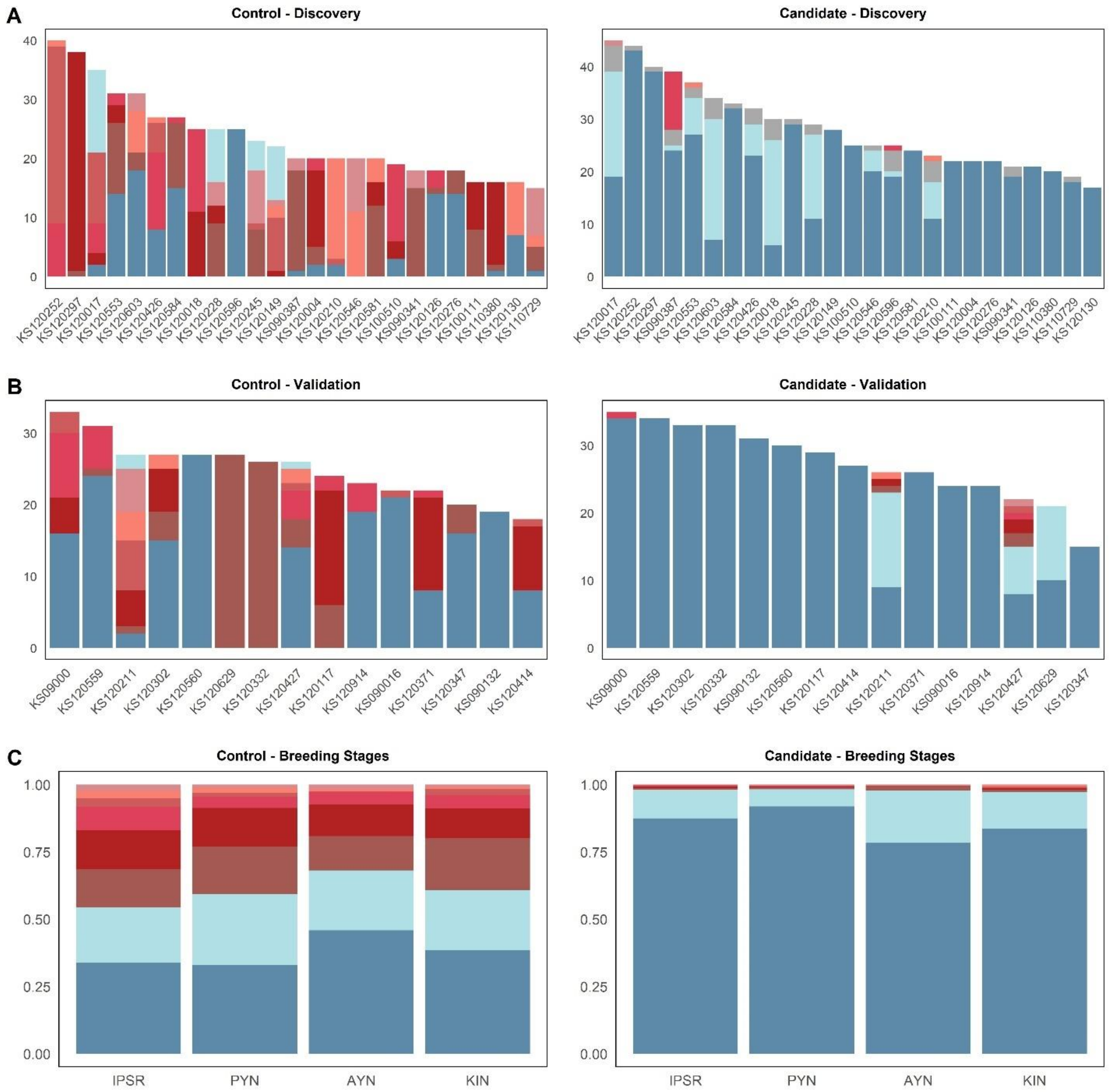


**Figure 19** Testing independent breeding panels for sign epistasis. Allele frequency count of the candidate three-way epistatic interaction involving segments of chromosomes **2D** (S2D_91221807), **3B** (S3B_713887174), and **4A** (S4A_693794455). **A** represents the data from the discovery populations. **B** displays data from the validation population. **C** consists of data from the last four stages of the breeding pipeline. In all plots, the control interaction, randomly selected and involving segments of chromosomes **1D** (S1D_463434894), **2B** (S2B_163607648) and **5A** (S5A_298295448), contrasts the candidate interaction with the null hypothesis.
